# Endosomal LRBA regulates the endo-lysosomal pathway

**DOI:** 10.1101/2024.02.07.579084

**Authors:** Viktória Szentgyörgyi, Leon Maximilian Lueck, Daan Overwijn, Nadia Zoeller, Maria Hondele, Anne Spang, Shahrzad Bakhtiar

**Author notes:** corresponding authors Correspondence: Shahrzad Bakhtiar MD PhD, ORCID 0000-0002-9898-1542 Goethe University Frankfurt University Hospital Department of Pediatrics Frankfurt am Main, Germany Anne Spang, ORCID:0000-0002-2387-6203 Biozentrum University of Basel Spitalstrasse 41 CH-4056 Basel Switzerland. equal contribution (Viktória Szentgyörgyi, Leon Maximilian Lueck).

## Abstract

Deleterious mutations in the *LRBA* (Lipopolysaccharide Responsive Beige-like Anchor protein) gene cause severe childhood immune dysregulation. The clinical manifestations of LRBA deficiency syndrome are highly variable. Thus, the complexity of the symptoms involving multiple organs and the broad range of unpredictable clinical manifestations complicate the choice of therapeutic interventions. Although LRBA has been linked to Rab11-dependent trafficking of the immune checkpoint protein CTLA-4, its precise cellular role remains elusive. We show that LRBA, however, does not colocalize with Rab11. Instead, LRBA is recruited by members of the small GTPase Arf protein family to the TGN and to Rab4^+^ endosomes, where it controls intracellular traffic. In patient-derived fibroblasts, loss of LRBA led to defects in the endosomal pathway yielding the accumulation of enlarged endolysosomes. Thus, LRBA appears to regulate flow through the endosomal system on Rab4^+^ endosomes. Our data strongly suggest functions of LRBA beyond CTLA-4 trafficking and provide a conceptual framework to develop new therapies for LRBA deficiency.

**Summary:** LRBA-deficient patients exhibit enlarged functional endolysosomes due to defects in recycling to the plasma membrane. LRBA localization on Rab4^+^ endosomes depends on Arf1 and Arf3, and is essential for a functional endosomal-lysosomal pathway. Our results could inform new treatment options.

## Introduction

LRBA (LPS-Responsive Beige-Like Anchor) belongs to the *be*ige *a*nd *Ch*ediak-Higashi (BEACH) domain containing protein family (Wang et al., 2001). Bi-allelic mutations in *LRBA* are associated with a severe immune deficiency and autoimmunity syndrome (Alangari et al., 2012; Lopez-Herrera et al., 2012). The average onset of the disease is at the age of two, and the only cure is allogeneic hematopoietic stem cell transplantation (alloHSCT) (Seidel et al., 2017; Bakhtiar et al., 2017; Tesch et al., 2020). The symptoms partially resemble that of CTLA-4 insufficiency (Lo et al., 2015). CTLA-4 is a plasma membrane receptor on regulatory T cells (Tregs) and suppresses autoimmune responses (Karandikar et al., 1996). In LRBA deficient patients, the overall and surface CTLA-4 protein levels are decreased due to its increased lysosomal degradation, which is otherwise prevented by LRBA. Consequently, LRBA deficiency is accompanied by a reduced number and suppressive capacity of Tregs. Based on the role of LRBA in CTLA-4 trafficking and on data on its *C.elegans* homolog SEL-2 (De Souza et al., 2007), LRBA has been linked to polarized endosomal recycling to the plasma membrane.

LRBA is a large, 319 kDa protein and contains an N-terminal concavalin-A like domain, a domain of unknown function (DUF), a protein kinase A binding motif (AKAP), the conserved BEACH domain, a non-canonical PH domain and C-terminal WD40 repeats, which are important for protein-protein interactions (Gebauer et al., 2004; Wang et al., 2001)(Fig. 1A). LRBA is expressed in most human tissues and was reported to be associated with the Golgi apparatus and vesicular structures . In support of this notion, the typical perinuclear localization of LRBA was lost when the Golgi was dispersed by brefeldin A (BFA) (Kurtenbach et al., 2017; Martinez-Jaramillo and Trujillo-Vargas, 2020). Since LRBA can co-immunoprecipitate with CTLA-4 and given that this interaction is abolished when the sorting motif in CTLA-4 tail is mutated, LRBA has been proposed to be a cargo adaptor (Lo et al., 2015). Yet, how LRBA would exert its function in the endosomal pathway, when the bulk of the protein is present on the Golgi remains elusive.

**Figure 1.**
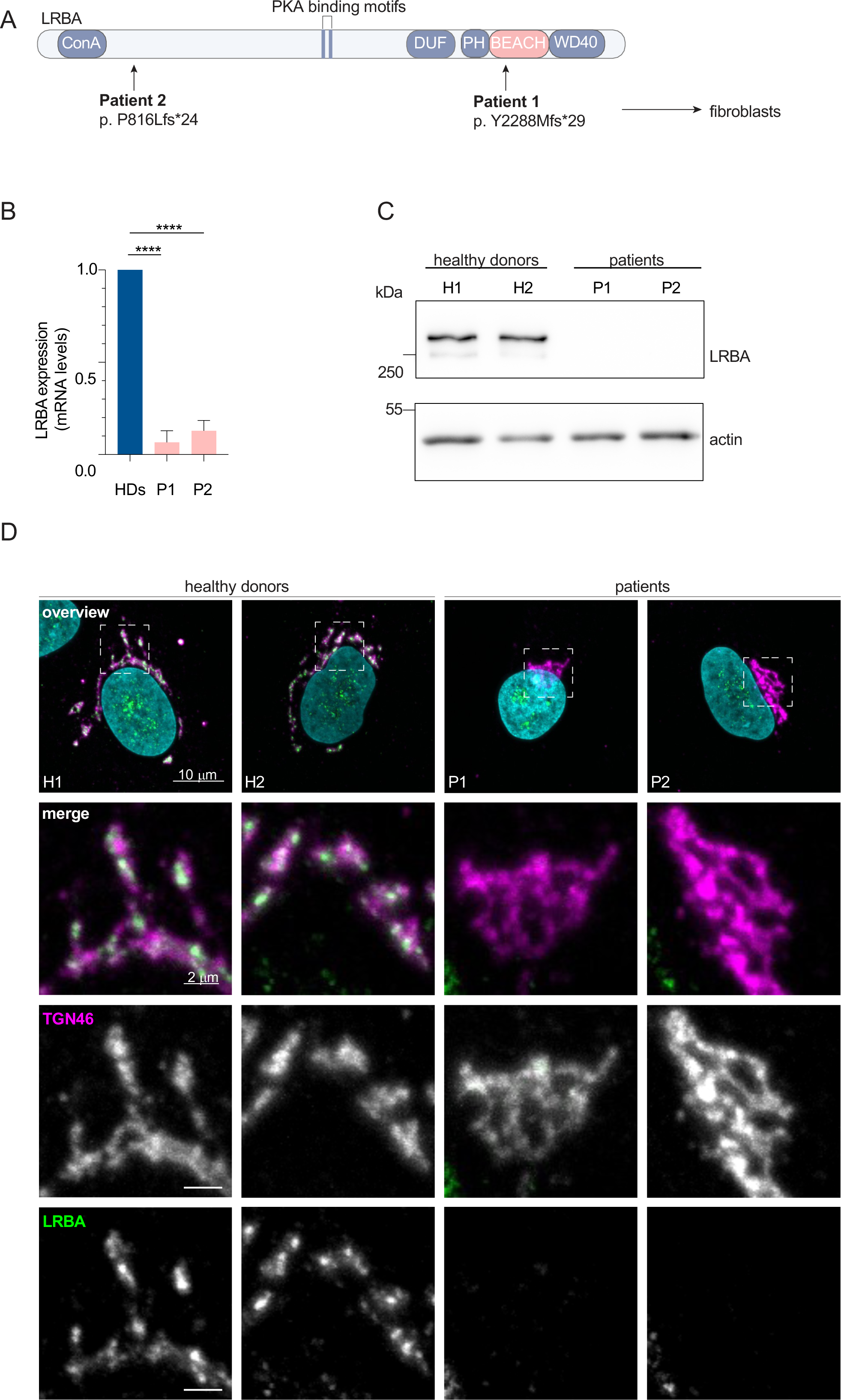
**Distinct point mutations in the *LRBA* gene of two LRBA deficient patients cause mRNA decay and loss of the protein**. **(A)** Schematic of LRBA protein structure with annotated domains. The genetic mutations carried by the two LRBA deficient patients investigated in this study are shown. Dermal fibroblasts obtained from these patients were used in this study. **(B)** LRBA mRNA levels in the two patient and two healthy donor (HD) fibroblast cell lines were determined by qRT-PCR. Mean and standard deviation are shown from n=4 biological replicates; one-way ANOVA using Dunnett’s multiple comparison, ****P<0.0001. **(C)** Immunoblot analysis of LRBA presence in fibroblasts of two HDs and two patient donors using polyclonal LRBA antibody and actin as a loading control. **(D)** Colocalization of LRBA and TGN46 in fibroblasts of HDs. LRBA is absent in patients’ fibroblasts. Cells were fixed, immunostained and imaged using a confocal microscope. Squares show magnification of the perinuclear area. The labeling of the single channels represents the color of the channel on the merged image. H1: HD 1, H2: HD 2, P1: patient 1, P2: patient 2.

Here, we report that in patient-derived dermal fibroblasts, the loss of LRBA leads to defects in endosome maturation accompanied by the accumulation of enlarged endolysosomes. We show that LRBA is recruited by members of the small GTPase Arf protein family to the trans-Golgi network (TGN) and by Arf1 and Arf3 to Rab4^+^ endosomes, where it controls intracellular trafficking. LRBA appears to play an important role in endolysosomal homeostasis. We propose that in LRBA deficiency the failure to correctly sort proteins for transport to the plasma membrane impedes with endosome- and endolysosome maturation. These findings expand our understanding of the underlying pathomechanism in the LRBA deficiency syndrome.

## Results

### LRBA is absent in patients with LRBA deficiency and affects Golgi organization on a global scale

To gain a better understanding of the molecular function of LRBA in the endosomal system, we analyzed dermal fibroblasts of two LRBA patients. The two patients have mutations at amino acid position 816 and 2288, respectively (Fig. 1 A). We analyzed the expression of LRBA in these cells and detected strongly reduced LRBA expression on both the mRNA (Fig. 1 B) and the protein level (Fig. 1 C), confirming the LRBA deficiency. Within the LRBA gene, another gene, MAB21L2 is nested (Tsang et al., 2009). Our qRT-PCR analysis indicated that MAB21L2 transcripts are still present in both patients, although with somewhat reduced levels as compared to the controls (Fig. S1 A). Thus, LRBA deficiency in these patients does not affect MAB21L2 expression. Next, we analyzed LRBA localization in fibroblasts. In cells, obtained from two healthy donors, LRBA localized to the trans-Golgi network (TGN), as observed previously (Wang et al., 2001; Lo et al., 2015; Kurtenbach et al., 2017)(Fig. 1 D). Consistent with the western blot results, LRBA was not detectable in patients’ cells by immunofluorescence (Fig. 1 D). The TGN appeared more compacted in LRBA patients’ cells as compared to those of healthy donors (Fig. 1 D and 2 A), indicating that LRBA may play a role in maintaining Golgi morphology.

To corroborate our findings, we determined the perinuclear distribution and the volume of the TGN (Fig. 2 B-E). While the volume of the TGN remained unchanged (Fig. 2 D), the TGN distribution around the nucleus was reduced in LRBA deficient cells (Fig. 2 B and C) and in parallel, we observed an increase in the mean fluorescence intensity of TGN46 (Fig. 2 E). These observations are consistent with a role of LRBA in maintaining Golgi distribution in the perinuclear region. Despite the changes in Golgi distribution around the nucleus, the Golgi ribbon and cisternal organization *per se* appeared to be intact in LRBA deficient cells as determined by electron microscopy (Fig. 2 F). Thus, it is likely that LRBA affects Golgi organization on the global but not on the local scale.

**Figure 2.**
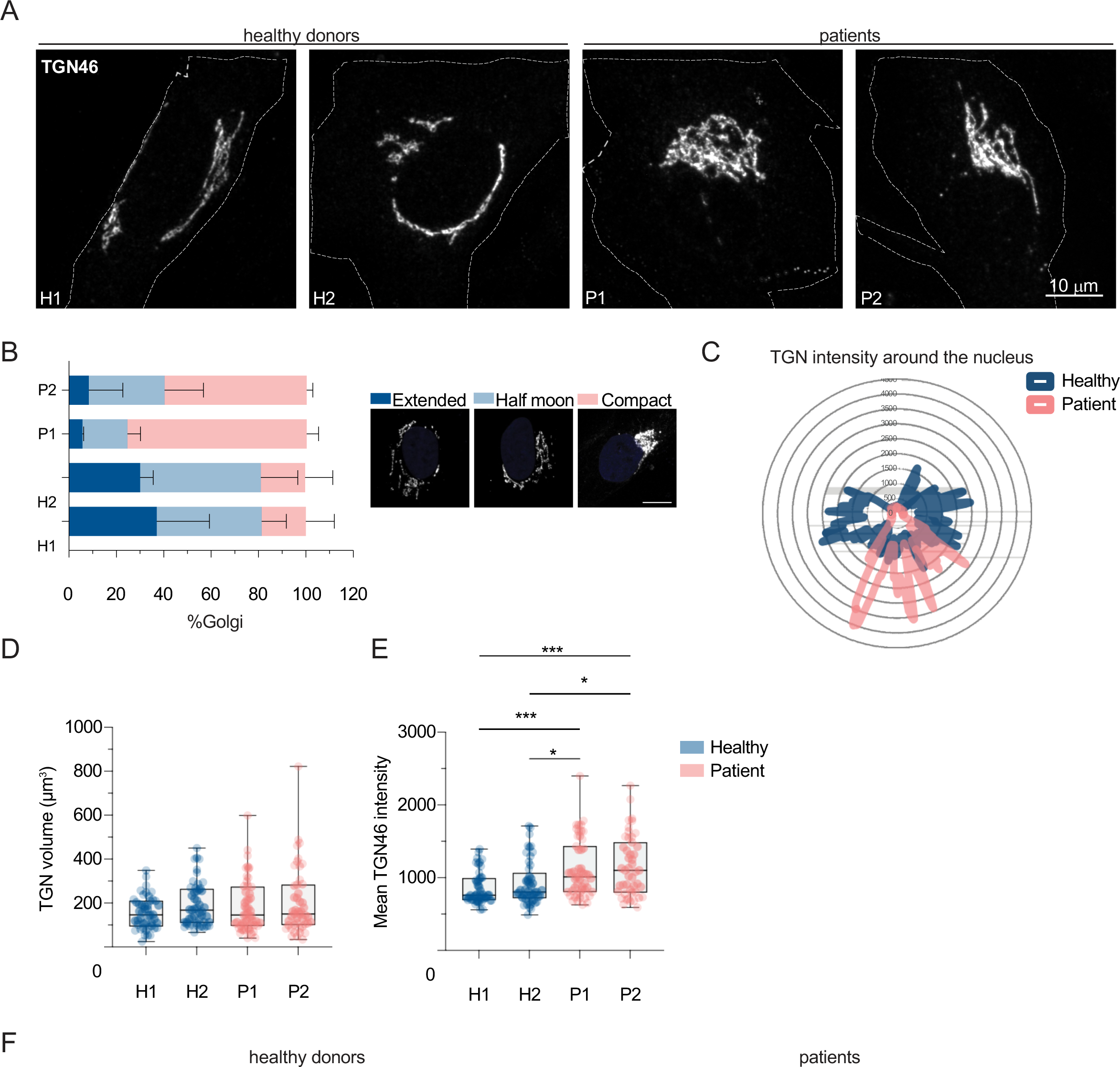
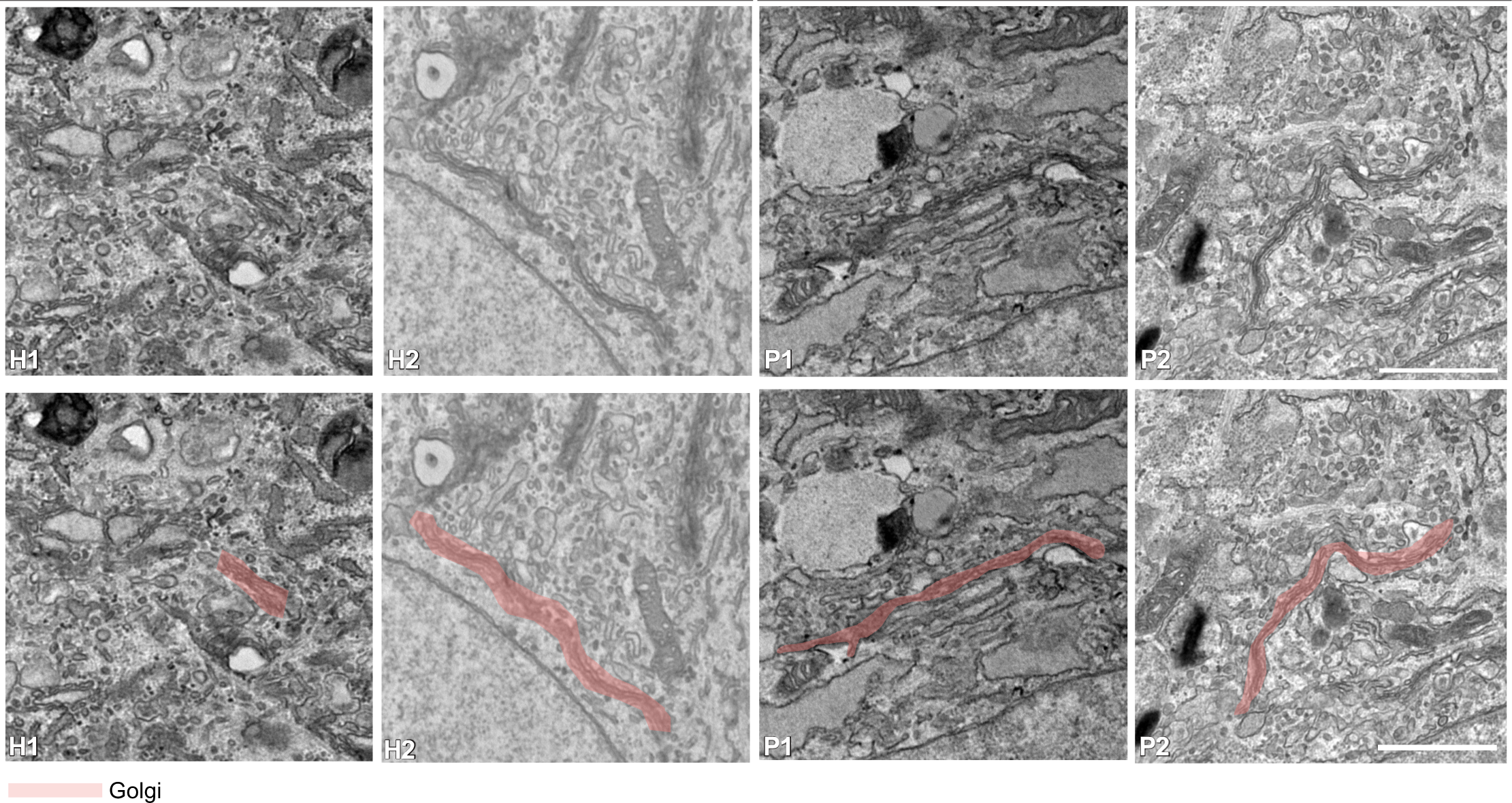
**TGN compaction in LRBA deficient patients’ cells**. **(A)** TGN morphology analysis of two HDs and two patient fibroblast cell lines visualized by the immunostaining of TGN46. Patients’ cells show compacted TGN morphology. Representative confocal images from n=3 biological replicates. Cell outlines are marked with a dashed line. H1: HD 1, H2: HD 2, P1: patient 1, P2: patient 2. **(B)** Measurements of TGN morphology (extended – dark blue, half moon – light blue, compact-pink) based on images taken in (A). Percentage of cells showing each category and standard deviation are shown; H1=47 cells, H2=47 cells, P1=53 cells, P2=49 cells from n=3 biological replicates. Legend shows a representative image for each morphology category. **(C)** Representative TGN46 signal distribution around the nucleus in H1 and P1 cells. **(D)** Quantification of TGN volume and **(E)** mean intensity of TGN46 signal based on images taken in (A). Mean and minimum to maximum are shown, box ranges from the first (Q1-25th percentiles) to the third quartile (Q3-75th percentiles) of the distribution. All data points are shown.; H1=46 cells, H2=49 cells, P1=51 cells, P2=47 cells from n=3 biological replicates; Kruskal-Wallis test using Dunnett’s multiple comparison, ***P=0.0007 (H1 vs. P1), ***P=0.0005 (H1 vs. P2), *P=0.0248 (H2 vs. P1), *P=0.0170 (H2 vs. P2) **(F)** TEM images of 2 HDs and two patients’ fibroblasts show intact Golgi cisternae. In the lower row, pink masks highlight Golgi stacks. Scale bar, 1 µm.

### Golgi-endosome trafficking is not impaired in the absence of LRBA

In order to assess whether the TGN compactness in LRBA deficient cells impacted trafficking from and to the Golgi, we analyzed the localization of the mannose 6-phosphate receptor (M6PR) (Fig. 3 A-F), which cycles between the TGN and endosomes in an adaptor protein complex-1 (AP1)- and retromer-dependent manner (Arighi et al., 2004). M6PR was not trapped in the TGN in patients’ cells (Fig. 3 A and B) and the colocalization of the receptor with the TGN (Fig. 3 A and C) was not changed between patients’ and healthy cells. The increase in the Mander’s coefficient of TGN overlap with M6PR in patients’ cells is most likely due to the more compact Golgi in those cells (Fig. 3 B). Most importantly, the trafficking of M6PR was not altered. To confirm these findings, we analyzed the colocalization of M6PR with the retromer subunit Vps35 (Fig. 3 D-F). We only observed very minor differences, which we consider unlikely to reflect a biological meaningful difference. Therefore, we conclude that the trafficking of M6PR is unaffected in LRBA deficient fibroblasts. Furthermore, we tested the recruitment of AP1 to the TGN (Fig. 3 G). AP1-positive carriers, which recycle cargo back from the recycling endosomes to the Golgi network (Hirst et al., 2012) were recruited normally in patient’s cells. Our data indicate that trafficking from and to the TGN is not altered in LRBA deficiency. Moreover, Golgi assembly after golgicide A (GCA) washout was not affected by the lack of LRBA (Fig. S1 B), indicating that LRBA is not essential for Golgi morphology establishment or overall function. Thus, even though at steady state, most of LRBA is at the Golgi, it does not seem to affect Golgi function locally.

**Figure 3.**
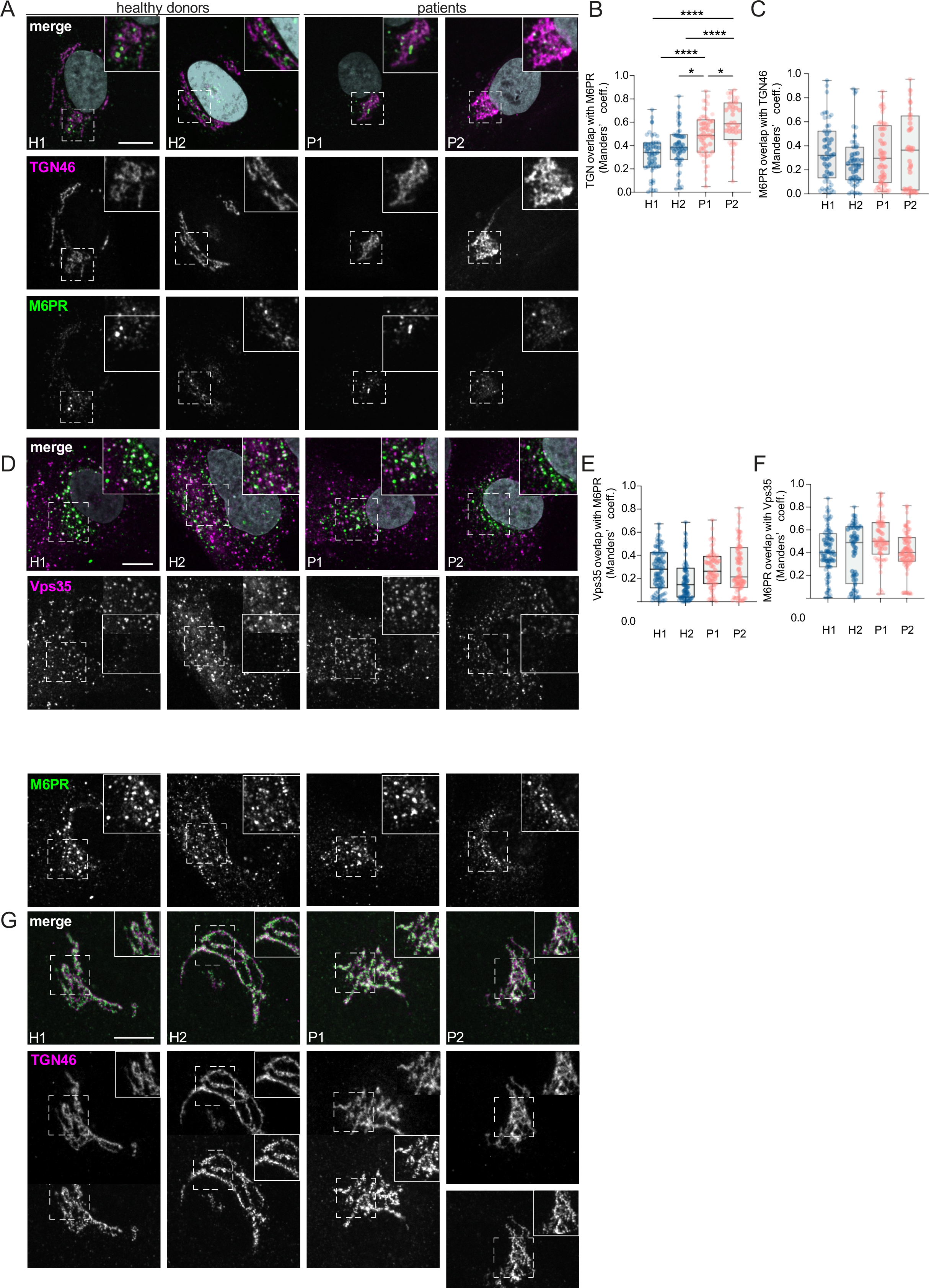
Golgi-endosome traffic is unimpaired in LRBA deficiency. (A) Colocalization analysis of TGN46 and M6PR in 2HDs and 2 patients’ fibroblast cell lines. Representative confocal immunofluorescence images of single focal planes. Squares show the magnified area. Inlays are shown in the top right corner of the images. The labeling of the single channels represents the color of the channel on the merged image. Scale bar, 10 μm. **(B-C)** Colocalization between TGN46 and M6PR was measured using Mander’s colocalization index. Mean and minimum to maximum are shown, box ranges from the first (Q1-25th percentiles) to the third quartile (Q3-75th percentiles) of the distribution. H1=47 cells, H2=47 cells, P1= 52 cells, P2=40 cells from n=3 biological replicates. **(B)** Mander’s coefficient of TGN46 overlap with M6PR. One-way ANOVA using Tukey’s multiple comparison, ****P<0.0001, *P= 0.0432 (H2 vs. P1), *P=0.0324 (P1 vs. P2). **(C)** Mander’s coefficient of M6PR overlap with TGN46. **(D)** Colocalization analysis of Vps35 and M6PR in 2HDs and 2 patient fibroblast cell lines. Representative confocal immunofluorescence images of single focal planes. Squares show the magnified area. Inlays are shown in the top right corner of the images. The labeling of the single channels represents the color of the channel on the merged image. Scale bar, 10 μm. **(E-F)** Colocalization between Vps35 and M6PR was measured using Mander’s colocalization index. Mean and minimum to maximum are shown, box ranges from the first (Q1-25th percentiles) to the third quartile (Q3-75th percentiles) of the distribution; H1=73 cells, H2=63 cells, P1= 56 cells, P2=68 cells from n=3 biological replicates. **(E)** Mander’s coefficient of Vps35 overlap with M6PR. **(F)** Mander’s coefficient of M6PR overlap with Vps35. **(G)** Immunofluorescence analysis of TGN46 and AP1 colocalization in 2HDs and 2 patient fibroblast cell lines. Representative confocal images of single focal planes. Squares show the magnified area. Inlays are shown in the top right corner of the images. The labeling of the single channels represents the color of the channel on the merged image. Scale bar, 10 μm.

### LRBA-loss leads to accumulation of enlarged endolysosomes in patient fibroblasts

The most striking and obvious phenotype of the patient-derived fibroblasts was a strong accumulation of enlarged endosomal/endolysosomal structures, which we observed by electron microscopy (Fig. 4 A). Endolysosomes are the fusion product of late endosomes (also dubbed multivesicular bodies, carrying cargo proteins for degradation) and lysosomes, and represent the active degradation compartment (Podinovskaia and Spang, 2018)(Fig. S2 A). In cells from healthy donors, endolysosomes appeared as electron-dense structures and contained many membrane layers as reported previously (Klumperman and Raposo, 2014)(Fig. 4 A, inlays, black arrowheads). In patient cells, late endosome-lysosome fusion could probably mostly still occur, but the electron-dense lysosome-derived material was segregated to one side in the enlarged endolysosome (Fig. 4 A, inlays, white arrowheads), indicating that either lysosomal function or endolysosomal maturation/lysosome reformation could be impaired. First, we confirmed that these accumulating structures are indeed endolysosomes. We stained the cells for the endolysosomal/lysosomal marker LAMP1 (CD107a) (Chen et al., 1985). Indeed, we observed a strong accumulation of LAMP1-positive structures in patient cells that were overall larger and more spread throughout the cell compared to cells from healthy donors (Fig. 4 B). Similarly, the vacuolar-ATPase accumulated on the enlarged endolysosomal structures (Fig. S2 B), and LAMP1 protein expression levels were increased in patient cells (Fig. S2 C). The altered gel mobility of LAMP1 suggested the presence of under-glycosylated protein species. Next, we asked whether the enlarged endolysosomes in patient cells would have retained their degradative capacity. Thus, we determined whether these endolysosomes were acidified and contained active proteases. To this end, we stained lysosomal compartments with LysoTracker, which showed a similar pattern as LAMP1 in patient cells, indicating proper acidification (Fig. S2 D). Similar results were obtained by cathepsin D immunostaining (Fig. S2 E). Cathepsins reach endolysosomes in an inactive pro-form, as so-called pro-cathepsins. Upon arrival in the endolysosome the pro-form is hydrolytically converted into the active form, which requires a low pH and is dependent on active proteases (Zaidi et al., 2008). Magic Red is a substrate that becomes fluorescent upon cleavage by cathepsin B and thereby allows the detection of cathepsin B activity. The Magic Red staining of catalytically active proteolytic structures was more spread and increased in intensity and in number (Fig. 4 C and D). In addition, we measured the levels of active, matured cathepsin D by western blot (Fig. 4 E). The levels of active cathepsin D were similar between healthy and patients’ cells (Fig. 4 E). Furthermore, the late endosome/endolysosomal marker Rab7 showed a similar pattern as LAMP1 (Fig. 4 F). Taken together, our data suggest that enlarged proteolytically active endolysosomes accumulate in LRBA deficient fibroblasts.

**Figure 4.**
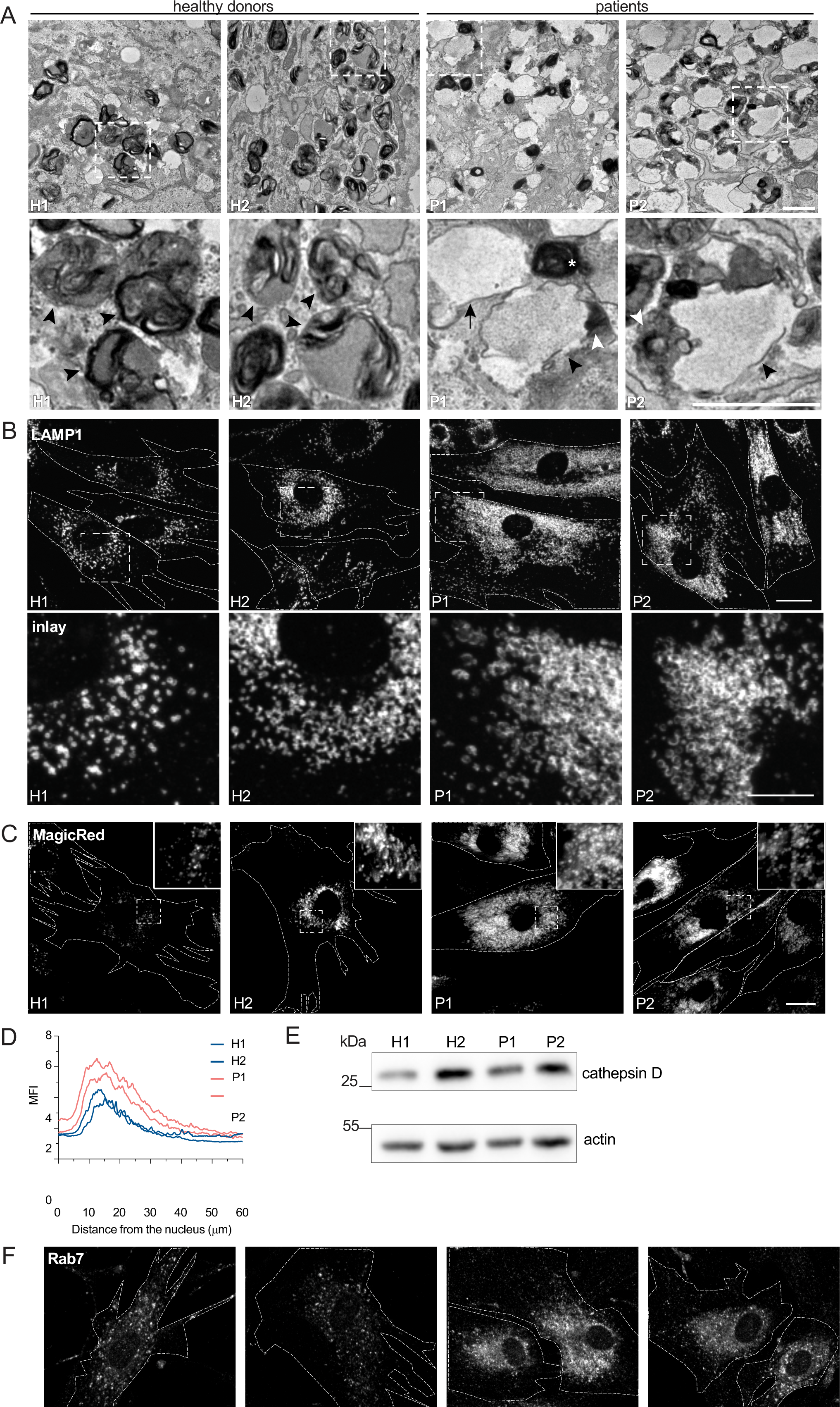
LRBA deficient fibroblasts show enlarged endolysosomes. (A) TEM analysis of accumulating endolysosomes in patients’ cells. Squares show magnification of the endolysosomal structures. HDs showed electron dense endolysosomes (black arrowheads). In contrast, in patients’ cells, endolysosomes (black arrowheads) showed restricted degradative (electron dense) domains (white arrowheads). Lysosomes (white star) and endosomes (black arrow) are shown. 2 HDs and 2 patient fibroblast lines were embedded and analyzed. Scale bar, 1 μm. **(B)** Immunofluorescence analysis of accumulating, enlarged (endo)lysosomes in patients’ cells. Fibroblasts were fixed with methanol and stained for LAMP1. Cell outlines are marked with a dashed line. Squares show magnification of the (endo)lysosomes. Representative confocal images are shown. Scale bar, 20 μm, inlays 10 μm. **(C)** Analysis of cathepsin B activity in LRBA deficient fibroblasts. LRBA deficient patients’ and healthy fibroblasts were plated onto imaging chambers and their lysosomes were visualized with Magic Red (indicating cathepsin B activity) and imaged live at 37°C, 5% CO_2_. Representative wide-field images are shown. Cell outlines are marked with a dashed line. Squares show the magnified area. Inlays are shown in the top right corner of the images. Scale bar, 20 μm. **(D)** Quantification of the MagicRed intensity along the nucleus-cell periphery axis. A line roi was drawn from the nucleus to the cell periphery and Magic Red intensity was measured. Values were normalized to the minimum of each cell and averaged per experiment. The mean of 3 biological replicates is plotted along the axis; H1= 45 cells, H2= 53 cells, P1= 51 cells, P2= 49 cells were analyzed. **(E)** Western blot analysis of matured cathepsin D (heavy chain) protein levels in healthy and patients’ fibroblasts using actin as a loading control. **(F)** Immunofluorescence analysis of Rab7^+^ late endosomes in 2 HDs and 2 patient fibroblast cell lines. Representative confocal images. Cell outlines are marked with a dashed line. Scale bar, 10μm.

### Receptor degradation is not impaired in LRBA deficient fibroblasts

If our assumption was correct, protein degradation in the enlarged endolysosomes in LRBA- deficient cells should still be functional. Thus, we followed the uptake and degradation of fluorescently labeled EGF, which upon binding to EGF receptors on the cell surface is endocytosed and subsequently degraded in endolysosomes (Tomas et al., 2014a). In cells from healthy donors, the EGF-TexasRed signal accumulated intracellularly within 15 min and was largely decayed after 60 min (Fig. 5 A and B). Even though the initial uptake of EGF within 15 minutes was lower in patients’ cells, EGF degradation occurred with similar kinetics to those of the healthy control confirming that the enlarged endolysosomes are still active. The lower intracellular EGF levels at 15 minutes of uptake could be explained with the reduced levels of EGF receptor observed in LRBA patient cells (Fig. S2 F). Yet, these data still did not explain why endolysosomes were bigger in patients’ cells. One possibility could be that the entire endosomal pathway would be affected, and already early endosomes would be enlarged. However, this does not appear to be the case because early endosomes were not consistently enlarged in the patient- derived cell lines. (Fig. 5 C and D), suggesting that the role of LRBA is downstream of early endosomes. To study later events of endosome maturation we tested EGFR signaling in LRBA patients’ cells (Fig. S3 A-C). EGF stimulation of EGF receptor on the cell surface induces autophosphorylation and tyrosine-kinase activity which stimulates signaling cascades in the cell. Activated EGFRs undergo rapid endocytosis and remain active on the surface of early endosomes (Tomas, Futter, and Eden 2014). In order to attenuate signaling, EGFRs need to be sorted into intraluminal vesicles (ILV) inside the endosome. Later, these multivesicular bodies/late endosomes fuse with lysosomes and their content is degraded. If ILV formation is perturbed, prolonged EGFR activation and downstream signaling can be observed (Eden et al., 2012). Thus, we stimulated LRBA patient and healthy cells with EGF and followed activated and total EGFR levels (Fig. S3 A and B). Again, we observed lower levels of EGFR in patient cells, yet the signaling pathway was activated, which we measured by blotting pEGFR and pERK (Fig. S3 A- C). The signaling attenuation and EGFR degradation kinetics were similar between healthy and patients’ cells. Thus, our data indicate that EGFR sorting into ILVs is not severely affected and that there is most likely a defect either during endosomal recycling or endolysosome-to-lysosome maturation.

**Figure 5.**
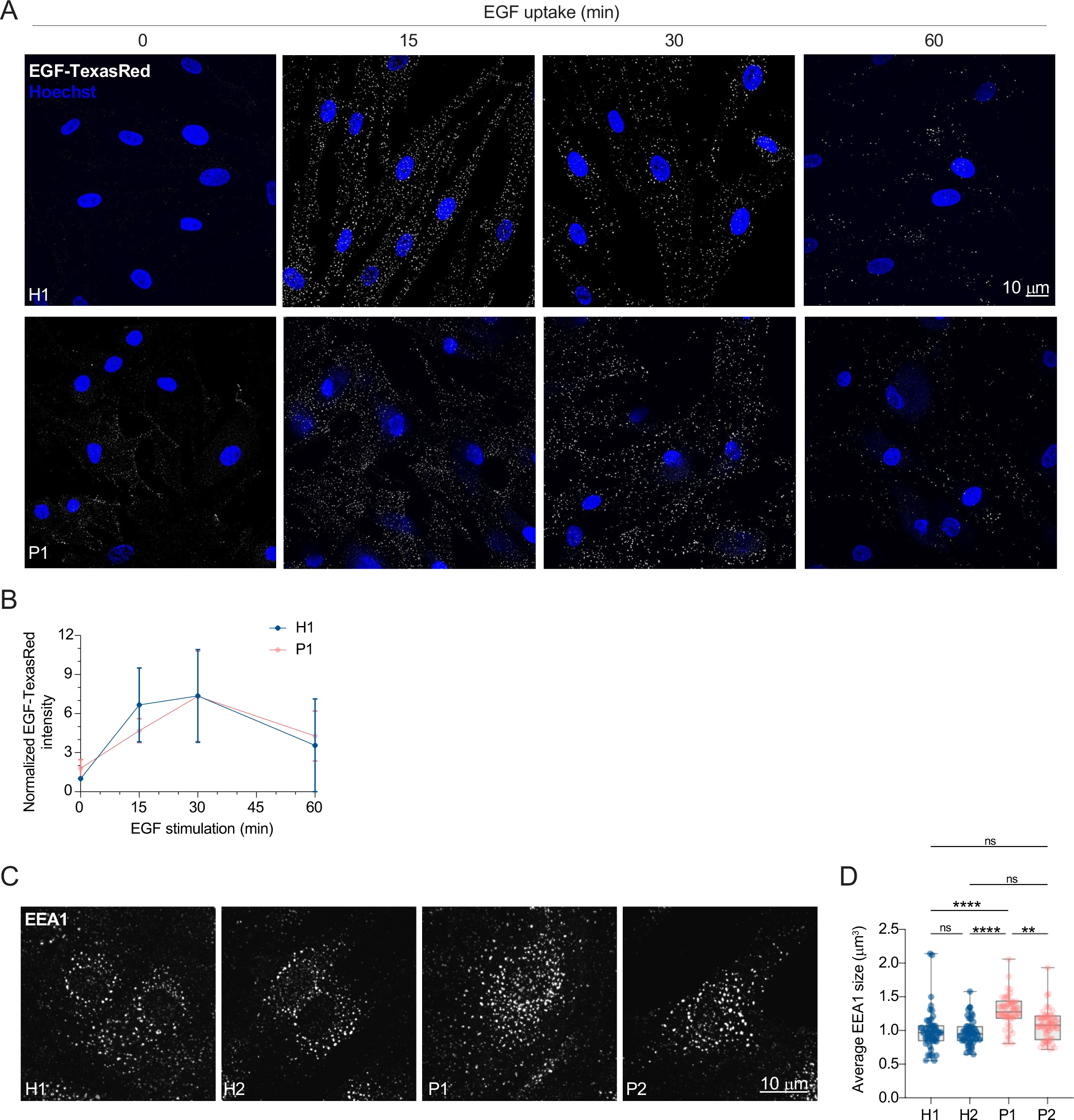
Lysosomal degradation is unimpaired in LRBA deficiency. (A) EGF-TexasRed uptake and degradation assay shows unimpaired degradation in LRBA deficiency. H1 and P1 fibroblast cells were serum starved for 3 hours and then were incubated on ice with EGF-TexasRed for 30 minutes. Cells were then washed 3x with ice-cold PBS and incubated with unlabeled EGF for indicated time points at 37°C. Cells were rinsed with ice-cold PBS, fixed with 4% PFA and mounted in the presence of Hoechst dye to visualize nuclei. Overview confocal images of EGF- TexasRed signal are shown for each time point. Maximum intensity projection of confocal Z-stacks. **(B)** Measurement of EGF-TexasRed fluorescence intensity based on images taken in (A). Integrated density of EGF-TexasRed per cell was measured, averaged and normalized to H1 levels at 0 time point for each experiment; H1(0min)= 38 cells, H1(15 min)= 58 cells, H1(30 min)= 67 cells, H1(60min)= 44 cells, P1(0min)= 36 cells, P1(15 min)=51 cells, P1(30 min)=45 cells, P1(60min)= 50 cells were analyzed from n=3 biological replicates. Mean and standard deviation is shown. **(C)** Immunofluorescence analysis of EEA1 positive early endosomes in healthy and LRBA patients’ fibroblasts. Maximum intensity projection of confocal Z-stacks. **(D)** The average size of early endosomes per cell was measured. Mean and minimum to maximum are shown, box ranges from the first (Q1-25th percentiles) to the third quartile (Q3-75th percentiles) of the distribution. All data points are shown. H1= 43 cells, H2= 44 cells, P1= 40 cells, P2= 40 cells were analyzed from n=3 independent experiments; Kruskal-Wallis test using Dunnett’s multiple comparison. ****P<0.0001, **P=0.0093.

### LRBA does not regulate the degranulation of CD8^+^ T or NK cells

Our data show that in LRBA-deficient fibroblast enlarged endolysosomes accumulate but still can degrade their content. Intriguingly, similar endolysosomal structures have been observed in Chediak−Higashi syndrome (CHS) patients’ cells from several cell types (Burkhardt et al. 1993; Stinchcombe, Page, and Griffiths 2000). The LYST protein, responsible for CHS disease, is a founding member of the BEACH domain protein family to which LRBA also belongs. This disease emerges from impaired cytotoxicity of natural killer (NK) and cytotoxic T cells (Abo et al., 1982; Baetz et al., 1995). These cells use lysosome-related lytic granules that contain, in addition to typical lysosomal proteins, perforin and granzymes (Fig. S3 D). The exocytosis of these granules at the cell-cell contact site – the immunological synapse - results in the induction of apoptosis in the target cell (Krzewski and Coligan, 2012). In CHS, although granule biogenesis is unimpaired, lytic granules are enlarged and fail to fuse with the plasma membrane (Stinchcombe et al., 2000; Gil-Krzewska et al., 2018). To test whether LRBA deficiency also leads to impaired exocytosis of lytic granules (degranulation) in CD8^+^ cytotoxic T cells and NK cells, we performed degranulation assays on patients’ cells (Fig. S3 E and Fig. S4). During the fusion of lytic granules with the plasma membrane, LAMP1 (CD107a) becomes exposed on the cell surface, which can be used as a marker for flow cytometric analysis of degranulating cells (Alter et al., 2004). T cells were activated with PMA and ionomycin (Fig S3 F, S4 A-C), while NK-cells were stimulated with interleukine-2 (IL-2) and K562 cells to cause surface exposure of LAMP1 (Fig. S4 D and E). The difference in degranulation was given by the ratio of mean LAMP1 fluorescence intensities (MFI) between patient and healthy donor (Fig. S4 C and E). As expected, under basal conditions, T cells did not show degranulation in either healthy donors or in the two LRBA patients (Fig. S4 A). Upon stimulation, degranulation of patient-derived CD8^+^ T cells and NK cells was almost as efficient as of cells from healthy donors. These results indicate that albeit the similarity in the accumulation of enlarged endolysosomes in CHS and in LRBA deficiency, LYST and LRBA have distinct cellular functions.

### LRBA is recruited by Arf1 and Arf3 onto Rab4+ endosomes

In order to gain more mechanistic insights into the role of LRBA in the endosomal trafficking, we switched to HeLa cells, which also endogenously express LRBA (Fig. S5 A and B). Similar to what we observed in fibroblasts, most of the LRBA accumulated at the TGN and colocalized with TGN46 and the small GTPase Arf1 but not with the cis-Golgi marker giantin (Fig. 6 A, S5 C). More importantly, we also detected LRBA on vesicular structures in the cell periphery (Fig. S5 B, inlay). We next aimed to uncover the identity of these structures by determining LRBA colocalization with different endosomal markers (Fig. 6 B). To our surprise, LRBA did not colocalize with Rab7^+^ late endosomes/endolysosomes or LAMP1^+^ lysosomes/endolysosomes. As expected, LRBA did not colocalize with M6PR either (Fig. S5 D). These data indicate that the observed endolyosomal enlargement in LRBA deficient fibroblasts might be a consequence of disturbances upstream in the pathway. LRBA appeared juxtaposed to Rab5^+^ and Rab11^+^ compartments; clearly on separate domains or structures from these markers (Fig. 6 B). However, we observed colocalization of LRBA with Rab4^+^ endosomes (Fig. 6 B), TGN46 and Arf1 in the cell periphery (Fig. 7 A). Consistent with this finding, LRBA has been reported to be sensitive to the ArfGEF inhibitor BFA (Kurtenbach et al., 2017; Martinez-Jaramillo and Trujillo-Vargas, 2020). Indeed, LRBA localization to the Golgi and the structures in the periphery was lost upon both BFA and GCA treatment (Fig. S5 E), indicating that Arf proteins are required for LRBA recruitment to Rab4^+^ endosomes. It has been shown previously that Arf1 and Arf3 are present on Rab4^+^ endosomes (D’Souza et al. 2014; Wong-Dilworth et al. 2023). To test whether Arf1 and/or Arf3 are required for LRBA recruitment, we used Arf1 knockout (KO), Arf3 KO and Arf1+3 double KO (dKO) HeLa cells (Pennauer et al., 2022) and stained them for endogenous LRBA (Fig. 7 B). While LRBA showed similar distribution in the single Arf1 KO and Arf3 KO cells as in the parental HeLa cell line, it was lost from endosomal structures in Arf1+3 dKO cells (Fig. 7 B and C). Intriguingly, in the dKO cells, LRBA localization to the Golgi was not affected. We assume that Arf4 and Arf5, which reside on the Golgi, could most likely compensate for the loss of Arf1 and 3 in the recruitment of LRBA to the Golgi.

**Figure 6.**
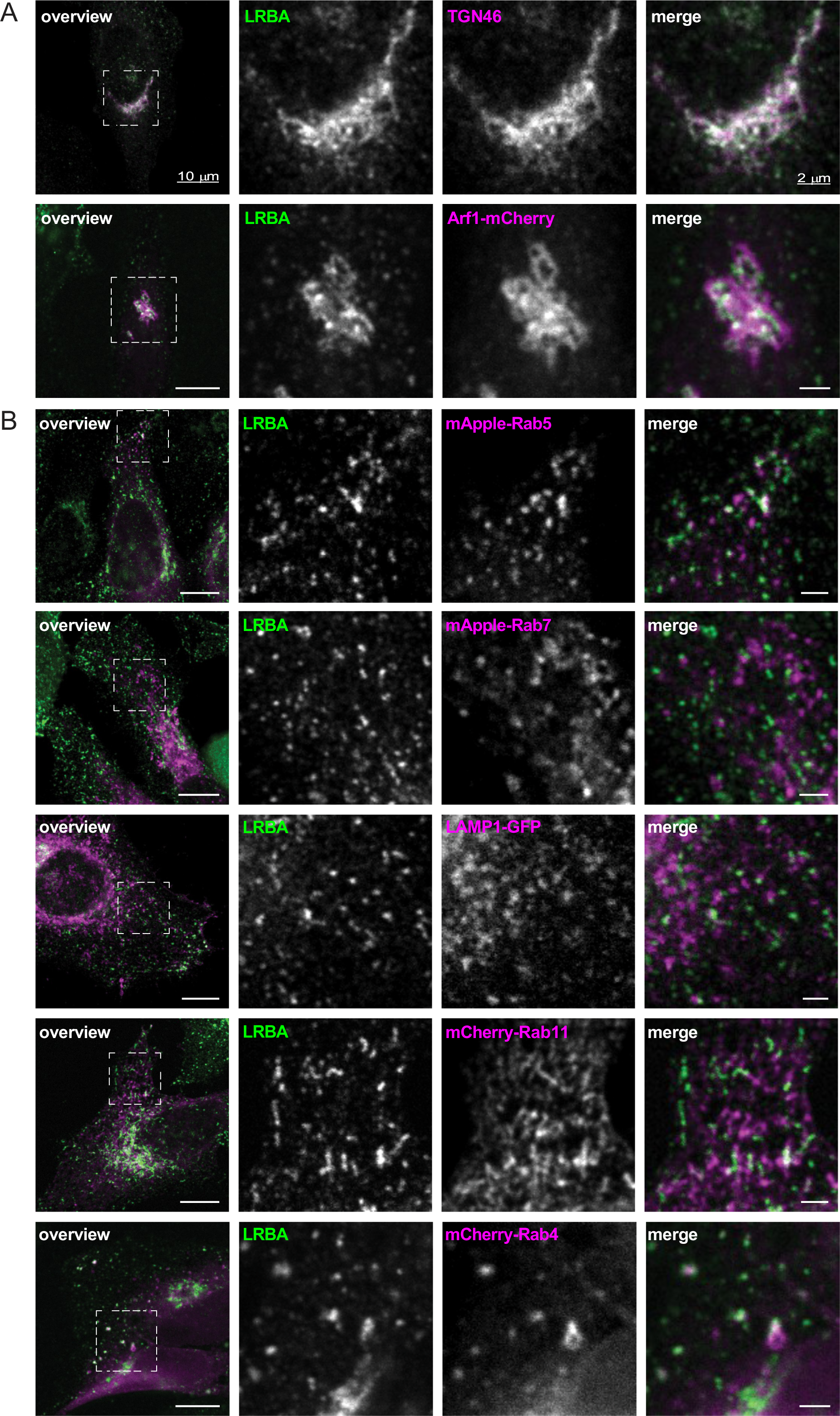
**LRBA colocalizes with the TGN and with Rab4^+^ endosomes in HeLa cells**. **(A)** Colocalization analysis of LRBA with TGN46 and Arf1 in HeLa cells. For the colocalization analysis with the TGN, HeLa cells were fixed with 4% PFA and stained for TGN46 and endogenous LRBA. For colocalization analysis with Arf1, HeLa cells were transfected with ARF1-mCherry, fixed with 4% PFA and stained for endogenous LRBA. Squares show magnification of the perinuclear area. The labeling of the single channels represents the color of the channel on the merged image. **(B)** Colocalization analysis of LRBA and different endosomal markers. LRBA colocalizes with Rab4 and found juxtaposition to Rab11 recycling endosomes and to Rab5 early endosomes. HeLa cells were transfected with mApple- Rab5, mApple-Rab7, LAMP1-GFP, mCherry-Rab11, mCherry-Rab4 respectively and stained for endogenous LRBA. Representative confocal images of single focal planes are shown. Squares show the magnified areas. The labeling of the single channels represents the color of the channel on the merged image. Scale bar, 10 μm, inlays 2 μm.

**Figure 7.**
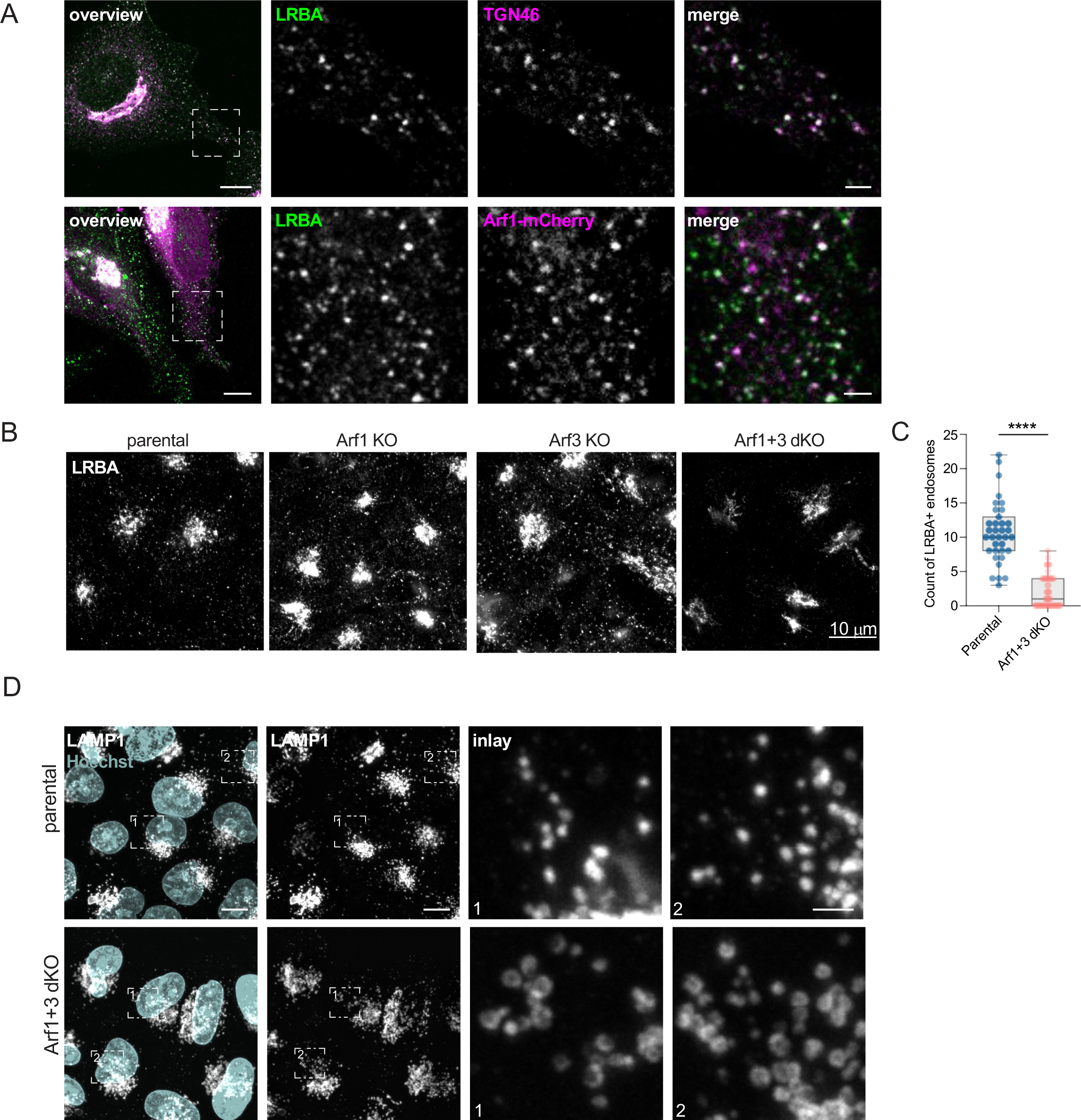
LRBA is recruited onto endosomes by Arf1 and Arf3. (A) Colocalization analysis of LRBA and TGN46 or Arf1 on endosomes in HeLa cells. For the colocalization analysis with the TGN, HeLa cells were fixed with 4% PFA and stained for endogenous TGN46 and LRBA. For colocalization analysis with Arf1, HeLa cells were transfected with ARF1-mCherry, fixed with 4% PFA and stained for endogenous LRBA. Squares show magnification of the perinuclear area. The labeling of the single channels represents the color of the channel on the merged image. Scale bar, 10 μm, inlays 2 μm. **(B)** LRBA is lost from endosomes in Arf1 and Arf3 dKO HeLa cells. Note that LRBA is still present on the Golgi. Parental, Arf1 KO, Arf3 KO, Arf1+3 dKO HeLa cells were seeded on coverslips, fixed, and stained for endogenous LRBA. Maximum intensity projections of confocal images are shown. **(C)** The number of LRBA^+^ endosomes per roi was measured. Mean and minimum to maximum are shown, box ranges from the first (Q1-25th percentiles) to the third quartile (Q3-75th percentiles) of the distribution. All data points are shown. Parental= 35 cells, Arf1+3 dKO= 34 cells were analyzed from n=3 independent experiments; unpaired two-tailed t test, ****P<0.0001. **(D)** (Endo)lysosomal structures are enlarged in Arf1+3 dKO cells. Parental and Arf1+3 dKO HeLa cells were seeded on coverslips, fixed, and stained for LAMP1 and with Hoechst. Squares show magnification of the (endo)lysosomes.

Our data indicate that Arf1 and Arf3 are required for recruitment of LRBA to Rab4^+^ endosomes and that loss of the endosomal LRBA pool is responsible for the accumulation of enlarged endolysosomes. Thus, we hypothesized that Arf1+3 dKO cells would also display enlarged endolysosomes. Indeed, LAMP1^+^ endolysosomes were enlarged in the absence of Arf1 and Arf3 (Fig. 7 D). To determine whether LRBA and Arf1/3 could interact directly, we turned to an *in silico* approach. We first predicted the structure of LRBA using Alphafold monomer© (Evans et al., 2021; Jumper et al., 2021)(Fig. 8 A and B). The predicted LRBA structure showed four α- solenoid regions (green), which are classical protein-protein interaction regions, connected by flexible linkers creating a rod-like structure. These α-solenoid regions contain the earlier described DUF domain (dark green). These regions are flanked on one side by several domains, which include concavalin-A like (cyan), PH (magenta), BEACH (red), and WD40 domain (yellow). Next we used Alphafold multimer © (Evans et al., 2021; Jumper et al., 2021) to predict potential interaction sites of LRBA with either Arf1 (Fig. 8 C) or Arf3. In most of our models, we observed that both Arf1 and Arf3 are predicted to potentially interact using conserved amino acid Ile^49^ (Fig. 8 C and D, S5 F) with two different hydrophobic pockets within LRBA solenoid regions around, either the Leu^861^ or Ile^918^, both of which are evolutionary conserved (Fig. 8 C and E). Additionally, several models showed Arf1 Asp^52^ interacting with LRBA Arg^910^. Thus, our *in silico* data revealed potential conserved binding sites between Arfs and LRBA, suggesting a direct interaction of Arf1 with LRBA.

**Figure 8.**
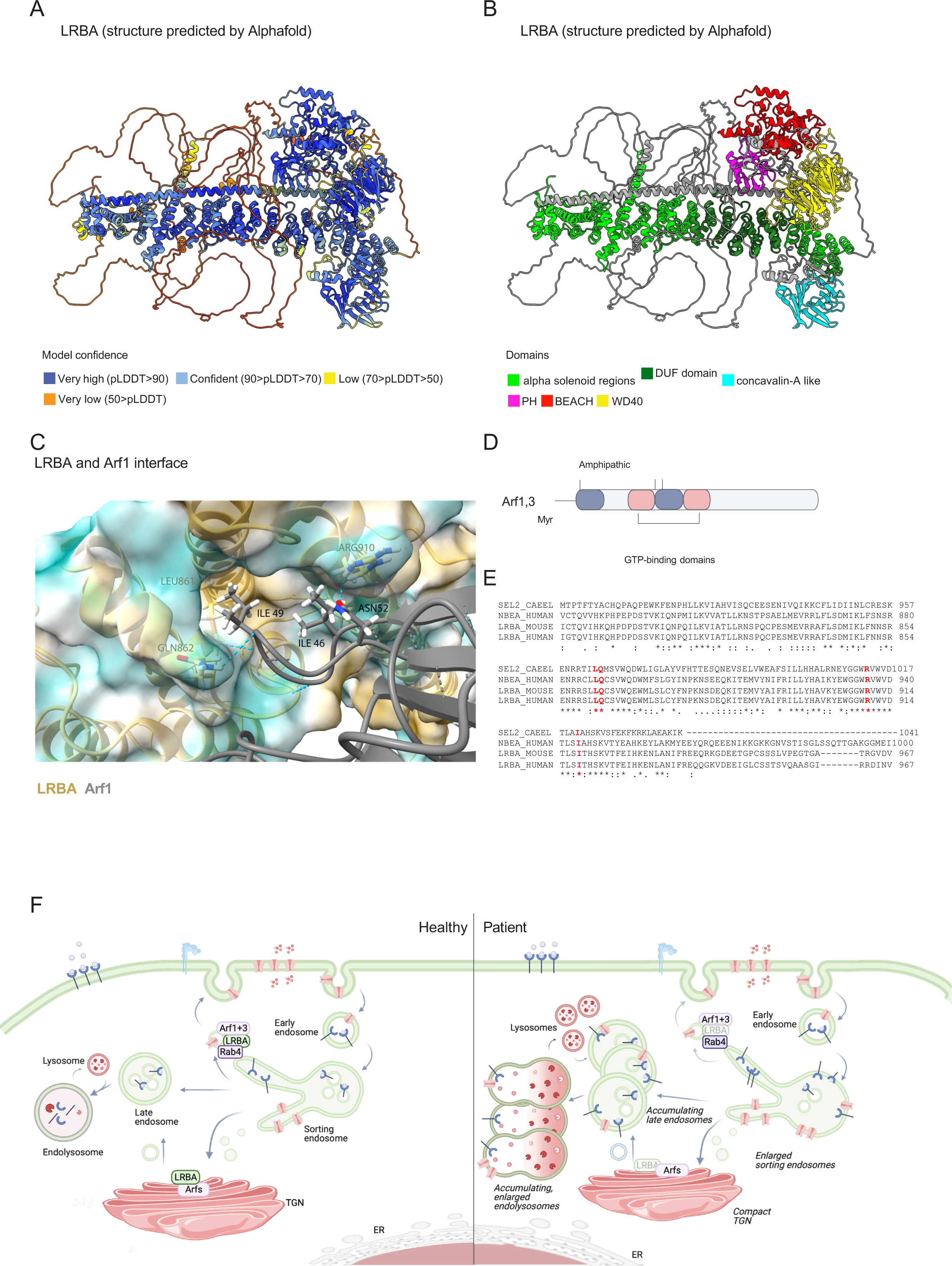
Structure of LRBA and potential interaction site with Arfs as predicted by Alphafold. (A-B) Alphafold prediction of human LRBA structure. Model confidence values **(A)** and domains **(B)** are indicated with colors. **(C)** Predicted interaction sites between LRBA and Arf1 using Alphafold monomer©. **(D)** Schematic of Arf1/3 structure and its domains. The conserved amino acid isoleucine 49 of Arf1/3 is shown since most of our models indicated to potentially interact with LRBA. **(E)** The aminoacids leucine 861, arginine 910 and isoleucine 918 of LRBA were predicted to interact with Arf1 and Arf3. All three amino acids are conserved across species. The amino acid sequences of the *C.elegans* SEL-2 (SEL2-CAEEL), the mouse and the human LRBA and the human neurobeachin (NBEA) were aligned. Labels: (*) conserved sequence; (:) conservative mutation; (.) semi-conservative mutation; (-) gap. **(F)** The role of LRBA in the endosomal system in health and disease.

## Discussion

In this study we show that LRBA is recruited to Rab4^+^ endosomes by Arf1 and Arf3 through potential direct interaction with these small GTPases. This LRBA localization seems to regulate the function of the endolysosomal pathway; when endosomal LRBA recruitment is perturbed, endolysosomes become enlarged. The endolysosomes are probably still functional when LRBA is lost because they were acidified, and transport from the TGN to the lysosomes and the maturation of lysosomal enzymes appeared to be functional. Thus, our data reveal a novel role of LRBA in the endosomal pathway (Fig. 8 F). Whether the enlarged endolysosomes, which appear to be defective in the lysosomal maturation process, are a direct consequence of LRBA deficiency or whether they are a consequence of the defects in endosomal recycling needs to be established. We currently favor a model in which the defects in sorting and transport to the plasma membrane at Rab4^+^ endosomes would prevent the endosome/endolyosomes to mature (Fig. 8 F). At least in T cells, recycling of CTLA-4 to the plasma membrane is LRBA-dependent (Lo et al., 2015). In addition, we observed reduced surface expression of EGFR in the absence of LRBA, consistent with a role of LRBA in recycling to the plasma membrane (Fig. 5 A and B, S2 F). Moreover, interfering with Rab11-dependent recycling to the plasma membrane yields functional, enlarged endolysosomes (Zulkefli et al., 2019).

Still, the bulk of LRBA was present on the Golgi. Moreover, we and others showed that LRBA becomes dispersed when the Golgi is abrogated by BFA and GCA (Kurtenbach et al., 2017; Martinez-Jaramillo and Trujillo-Vargas, 2020). Consistent with the Golgi localization of LRBA, the Golgi morphology was altered in LRBA deficient cells and became more compact. While we could not detect any trafficking defects, under-glycosylated LAMP1 became more prominent under these conditions, indicating a potential role for LRBA at the Golgi and that Golgi function might still be somewhat perturbed. In addition, LRBA was reported to be on Rab11^+^ recycling endosomes in T cells and renal collecting duct tissue (Lo et al. 2015; Yanagawa et al. 2023). When we revisited this colocalization, we only observed a slight overlap; LRBA was found to be juxtaposed to Rab11^+^ domains in HeLa cells. Instead, we found LRBA on Rab4^+^ endosomes. While it is conceivable that LRBA might be on different endosomes depending on the cell type, we favor the explanation that because of the greater spatial resolution in HeLa cells, we are able to separate LRBA from Rab11^+^ domains. Most importantly, we showed that the specific loss of the endosomal pool of LRBA is responsible for the major phenotype observed in patient cells.

We further established that LRBA is recruited to endosomes specifically by Arf1 and Arf3. We propose that this recruitment is mediated via a direct interaction between Arf1/3 and LRBA through the Ile^49^ site in the hydrophobic pocket of Arfs. Supporting our model, this hydrophobic pocket (Ile^49^,Gly^50^,Val^68^,Gly^69^,Ile^74^,Leu^77^) together with three other amino acids referred to as hydrophobic triad (Phe^51^, Trp^66^, Tyr^81^) build a hydrophobic surface which is often recognized by Arf effectors like ARHGAP21, COPI and GGA (Jacques et al., 2002; Sun et al., 2007; Ménétrey et al., 2007). This hydrophobic pocket is only exposed in the GTP-bound form and despite high variability in the structures of effectors, a hydrophobic residue, usually Ile or Leu from the effector interacts with the pocket (Chavrier and Ménétrey, 2010). Similarly, our Alphafold prediction indicates the LRBA residues Leu^861^ and Ile ^918^ for interaction with Arf1 and Arf3, respectively.

Besides the most prominent and classical localization on the Golgi, Arf1 and Arf3 were found on different endosomal membranes, such as Rab4^+^ endosomes (D’Souza et al., 2014) and on TGN46 positive Golgi derived vesicles (Boutry et al. 2023; Wong-Dilworth et al. 2023). Interestingly, we found LRBA on both Rab4^+^ and on TGN46 and Arf1 positive structures where LRBA regulates endosomal recycling. LRBA is also likely binding to Arf4 and/or Arf5, because in the Arf1/3 dKO, LRBA Golgi localization was unaffected. Moreover, the binding pocket is conserved among all these Arfs. The most prominent phenotype, we observed in LRBA deficient cells was the accumulation of enlarged endolysosomes, which is caused by the loss of the endosomal LRBA pool. Interestingly, similar findings have been reported in Chediak Higashi syndrome (CHS)(Burkhardt et al., 1993; Stinchcombe et al., 2000; Bowman et al., 2019). In the CHS disease giant lysosomes and lysosome related organelles like melanosomes are present in patients’ cells due to aberrant function of lysosome related organelles. Similarly to CHS, we find that LRBA does not affect lysosomal biogenesis and degradation and does not colocalize with LAMP1 under steady state conditions. Lysosomes were correctly acidified and contained catalytically active hydrolases like cathepsin D and B based on our Magic Red assay and western blot analysis. Degradation of EGF-TexasRed also occurred with similar kinetics as in control cells. In contrast to CHS disease, however, we could not detect any significant impairment in degranulation in patients’ CD8^+^ T cells and in NK cells, despite the observation that patients’ cells stayed slightly below the range of healthy controls. Based on our current data, we believe that LRBA does not result in the same level of degranulation deficiency as seen in other BEACH related diseases, such as CHS or Grey Platelet syndrome caused by mutations in NBEAL2 (Sowerby et al., 2017).

While lysosomal dysfunction was described for a plethora of diseases among which are SLE (Wang and Muller, 2015), Sjörgen disease (Sohar et al., 2005), Crohn’s disease (Lassen et al., 2016) and rheumatoid arthritis (Ansari et al., 2021), the enlarged endolysosomes in CHS and LRBA are functional in terms of protein degradation. Enlarged endosomes/endolyosomes appear to be a common feature of BEACH related diseases. It is tempting to speculate that the defect in α-granule precusor biogenesis, which is the underlying cause of Grey Platelet syndrome in NBEAL2 deficiency (Lo et al., 2018) is due to the enlarged endosomes/endolyosomes that also fail to mature into α-granule precursors. Thus, it is plausible that the underlying cause of BEACH related diseases is the inability to form functional lysosome-related organelles (LROs); such as cytolytic granules in CHS and α-granules in NBEAL2 deficiency (Bowman et al., 2019). Which LRO cannot be formed in LRBA-deficiency remains elusive. Moreover, how the observed perturbed endosomal traffic and the endolysosome accumulation lead to the manifestation of LRBA deficiency syndrome needs further investigation. It will be important to determine how LRBA regulates sorting on Rab4^+^ endosomes. Our study highlights the importance of improving transport through the endosomal system in the search for therapeutic options in LRBA-deficiency.

## Material and Methods

### Patient material

Informed consent was available according to the approved protocols from local institutional review board of Goethe University Frankfurt, Germany (IRB # 436/16). Skin fibroblast and peripheral blood samples were analyzed. Patient 1 is currently a 26 year-old female of Egyptian descent whose disease aggravated over the past years needing an alloHSCT at the age of 24 years, which was successful. Patient 2 is currently a 5 year-old boy of Libyan descent who developed severe early onset inflammatory bowel disease, also necessitating alloHSCT. Both patients had received abatacept prior to alloHSCT. The clinical course of Patient 1 has been previously published by our group (Bakhtiar et al., 2016). Patients 3 and 4 are 14- and 17- year-old non- transplanted patients under abatacept treatment, their blood samples were analyzed for degranulation assays.

### Cell culture

Dermal fibroblasts were isolated from skin biopsies of LRBA deficient patients 1 and 2 by explant culture and healthy donors according to the protocol (Zöller et al., 2008). The fibroblasts were cultured in high-glucose Dulbecco’s modified Eagle’s medium high glucose (4.5g/l) (Sigma- Aldrich) with 5% (v/v) fetal bovine serum (FBS), 100 U ml−1 penicillin G and 100 ng ml−1 streptomycin, at 37 °C and 7.5% CO_2_.

*ARF1, ARF3* and *ARF1+3* double KO HeLaα cells were established, mycoplasma tested and described elsewhere (Pennauer et al., 2022). HeLa CCL2 cells were kindly provided by Dr. Martin Spiess, with its identity authenticated by STR analysis by Microsynth AG (Balgach, Switzerland).

LRBA KO, wild type HeLa CCL2, *ARF1, ARF3* and *ARF1+3* double KO HeLaα cells and HeLa Flp-In-T-REx 3xFlag-EGFP-LRBA were grown in high-glucose Dulbecco’s modified Eagle’s medium (Sigma-Aldrich) with 10% fetal bovine serum (FBS, Biowest), 2 mM L-glutamine, 100 U ml−1 penicillin G and 100 ng ml−1 streptomycin, 1 mM sodium pyruvate (referred to as CM) at 37 °C and 5% CO_2_.

Expression of 3xFlag-EGFP-LRBA was induced with 1μg/mL tetracycline in the HeLa Flp-In-T- REx 3xFlag-EGFP-LRBA cell line (kindly provided by Dr. Serhiy Pankiv and Prof. Anne G. Simonsen, University of Oslo, Norway).

Heparin-anticoagulated blood samples were obtained from patients and healthy donors and peripheral mononuclear blood cells (PBMCs) were isolated from whole blood within 24 hours. Briefly, heparin-blood was diluted 1:2 with Dulbecco’s Phosphate-Buffered Saline (DPBS), layered on cell separation solution medium (Bicoll, Biochrom, Berlin, Germany) following centrifugation for 20 minutes at 800*g* without deceleration. After removing the top layer, the mononuclear cell containing ring was collected and washed twice with DPBS. The number of isolated PBMCs was determined using a cell counter (DxH500, Beckman Coulter, Krefeld Germany). Cells were cultured in RPMI 1640 medium (Gibco) supplemented with 10% FBS.

### CRISPR-Cas9 KO cell line generation

For CRISPR–Cas9-mediated KO, guide RNAs were selected using the GenScript© CRISPR design tool. Two guide RNAs were designed from two different exons for the target LRBA gene, gRNA against exon 22: CCATGCAGTCAAATATGAGT, gRNA against exon 38: GGTTACGCACAAATCGTCGC. Annealed oligonucleotides were cloned into Px458-GFP vector and Px459-Puro vector using the BbsI cloning site. In brief, 7.5x10^5^ HeLa cells were seeded per 10-cm dish. The following day, cells were transfected with 6 μg of the plasmids (3-3 μg for both targeted exon of LRBA) with Helix IN transfection reagent (OZ Biosciences). For control cells control vectors without gRNA insert were transfected. For selection, cells were treated with 1.5 μg/ml puromycin for 24 h before FACS sorting (for GFP^+^ cells). FACS sorting was carried out 48 hours after transfection. Cells were trypsinized and resuspended in cell-sorting medium (2% FCS and 2.5 mM EDTA in PBS) and sorted on a BD FACS Aria Fusion Cell Sorter.

GFP^+^ cells were collected and seeded onto 96 well plate. Cells were then expanded and LRBA expression was determined with western blot.

### Antibodies

The following antibodies were used in this study: sheep polyclonal anti-TGN46 (#AHP500G, BioRad, 1:2000), rabbit polyclonal LRBA antibody (#HPA023567, Atlas Antibodies, 1:1000), mouse AP1 100/3 hybridoma antibody (home-made), rabbit monoclonal EEA1 (C45B10) antibody (#3288, Cell Signaling, 1:2000), rabbit monoclonal anti LAMP1 (D2D11) XP® antibody (#9091, Cell Signaling, 1:200 for immunofluorescence, 1:1000 for western blot), rabbit monoclonal Rab7 (D95F2) XP® antibody (#9367, Cell Signaling, 1:400 for immunofluorescence, 1:1000 for western blotting), mouse monoclonal anti M6PR (cation dependent) (2G11) antibody (#ab2733, Abcam), rabbit polyclonal anti giantin antibody (#924302, BioLegend, 1:500), goat polyclonal anti Vps35 antibody (#ab10099, Abcam, 1:50), rabbit monoclonal anti EGF receptor (D38B1) XP® antibody (#4267, Cell Signaling, 1:1000), rabbit monoclonal phospho-EGF receptor (Tyr1068) (D7A5) XP® antibody (#3777, Cell Signaling, 1:1000), mouse monoclonal anti p44/42 MAPK (Erk1/2) (L334F10) antibody (#4696, Cell Signaling, 1:2000), rabbit polyclonal anti phospho-p44/42 MAPK (Erk1/2) (Thr202/Tyr204) antibody (#9101, Cell Signaling, 1:1000), mouse anti calnexin antibody clone 37 (#610523, BD Biosciences, 1:2000), mouse monoclonal anti Pan Actin (#LCU9001, Linaris, 1:1000), mouse monoclonal anti TCIRG1/ lysosomal ATPase V0 subunit a3 antibody (M01), clone 6H3 (#H00010312-M01, Abnova), rabbit monoclonal recombinant anti Cathepsin D [EPR3057Y] antibody (#ab75852, Abcam, 1:100 for immunofluorescence, 1:2000 for western blot), Hoechst 33342 Fluorescent Nucleic Acid Stain (#639, ImmunoChemistry Technologies, 1:200), goat anti-rabbit IgG(H+L) secondary antibody, AlexaFluor^TM^488 (#A-11034, Invitrogen, 1:500), goat anti-Mouse IgG (H+L) secondary antibody, AlexaFluor^TM^488 (#A11001, Invitrogen, 1:500), goat anti-mouse IgG (H+L) secondary antibody, AlexaFluor^TM^633 (#A-21052, Invitrogen, 1:500), donkey anti-goat IgG (H+L) secondary antibody, AlexaFluor^TM^488 (#A11055, Invitrogen, 1:500), donkey anti-mouse IgG (H+L) Secondary antibody, AlexaFluor^TM^568 (#A10037, Invitrogen, 1:500), donkey anti-sheep IgG (H+L) secondary antibody, AlexaFluor^TM^568 (#A21099, Invitrogen, 1:500), mouse monoclonal alpha-tubulin antibody (#T5168, Sigma-Aldrich, 1:10.000), goat anti-mouse IgG, (H+L), HRP-coupled (#31430, Pierce/Invitrogen, 1:10000), goat anti-rabbit IgG, (H+L), HRP- coupled (#31460, Pierce/Invitrogen, 1:10000), anti-CD107a-PB (#B13978, Beckman Coulter), anti-CD45-KrO (#B36294, Beckman Coulter), anti-CD3-APC (#IM2467, Beckman Coulter), anti-CD8-APC-AF700 (#B49181, Beckman Coulter), anti-CD69-PE (#IM1943U, Beckman Coulter), CD56-PC7 (#A21692, Beckman Coulter).

### DNA and plasmid sources

The following commercially available plasmids were obtained: mApple-Rab5a (Addgene #54944), mApple-Rab7a (Addgene #54945), LAMP1-GFP (Addgene #34831), mCherry-Rab11a (Addgene #55124, mCherry-Rab4a (Addgene #55125), pSpCas9(BB)–2A-GFP (pX458) (Addgene #48138), pSpCas9(BB)–2A-puro (pX459) (Addgene, #48139).

Arf1-mCherry was cloned by replacing EGFP in the pEGFP-N1-Arf1 (Addgene #39554) plasmid with mCherry. mCherry was amplified by PCR from the mCherry-Rab4a (Addgene #55125) plasmid and inserted with NEBuilder HiFi Assembly cloning kit (#E5520S, New England Biolabs).

### TEM

Fibroblasts were grown to 90% confluency on fibronectin (Sigma Aldrich) coated 18 mm round glass coverslips. Cells were fixed by adding a warm double strength fixative solution (5% glutaraldehyde (GA, Electron Microscopy Sciences, #16310) and 4% paraformaldehyde (PFA, Electron Microscopy Sciences, #15710) in 0.1M PIPES buffer (Sigma, #P6757 supplemented with 2mM CaCl_2_, pH 7-7.3)) to the culture medium into the dish (ratio 1+1) and mix very gently and incubated for 15 minutes at room temperature. Then the fixative-medium mixture was replaced by fresh single strength fixative solution (2% PFA, 2.5% GA in 0.1M PIPES) and incubated for 2 hours at RT and for 16 hours at 4°C. Then cells were washed 3 times for 10 minutes with cold 0.1M PIPES buffer. Fixed cells were rinsed first in PIPES buffer and then once in cacodylate buffer (0.1 M, pH 7.3) for 10 min. After two additional washes in cacodylate buffer, cells were post-fixed in 1% osmium tetroxide (Electron Microscopy Sciences, #19100), 0.8% potassium ferracyanide in 0.1M cacodylate buffer for 1 hour at 4°C. Coverslips were rinsed several times in cacodylate buffer and ultrapure distilled water then, *en block* stained with 1% aqueous uranyl acetate for 1 hour at 4°C in the dark. After several wash steps in ultrapure distilled water, cells were dehydrated in an ethanol series (30, 50, 75, 95% and 100%, Electron Microscopy Sciences, #15056) at 4°C followed by three additional changes of absolute ethanol. Samples were washed in acetone (Electron Microscopy Sciences, #15056) and finally embedded in a mixture of resin/acetone first and then in pure Epon 812 resin (Electron Microscopy Sciences, #14120) overnight. Coverslips were placed cell-side down on BEEM capsules (Electron Microscopy Sciences, #70010-B) filled with EPON. Polymerization was carried out for 48 hours at 60°C. After complete polymerization, coverslips were removed from EPON block using the nitrogen-hot water method. During the removal of the coverslip from the EPON block cells are transferred from the coverslip to the block surface. 70 nm thin serial sections were cut with diamond knife and placed on formvar-carbon coated copper grids, stained with uranyl acetate (Electron Microscopy Sciences, #22400) and Reynolds’s lead citrate and observed into a FEI Tecnai G2 Spirit Transmission Electron Microscope (TEM) operating at 80kV. Images were acquired using an EMSIS Veleta camera (top, side-mounted). The Camera operates using the RADIUS software from EMSIS.

### Immunostaining

HeLa cells were plated onto coverslips 24 hours prior to fixation. When indicated, transfected, cell fixation was performed 48 hours after cell plating. Cells were fixed in 4% paraformaldehyde, permeabilized with 0.1 % Triton X-100, blocked in PBS containing 5% FBS, and stained with the indicated primary antibodies followed by AlexaFluor conjugated secondary antibodies. Coverslips were mounted onto glass slides with Fluoromount G (Southern Biotech) or Vectashield and sealed with nail polish.

### Microscopy

Confocal images were acquired with Olympus Fluoview FV3000 system, using an UPLSAPO 60x/1.30 objective with silicone oil. Sampling speed was 8.0 μs/pixel. All images are representative from at least three independent sets of experiments. All images for corresponding experiments were processed with the same settings to ensure comparable results.

### Image analysis

Golgi morphology analysis was based on confocal Z-stack images of TGN46. Different Golgi phenotypes were categorized according to their extension around the nucleus (extended, half moon, compact).

For the TGN radar plots, two-pixel thick segmented lines with spline fit were drawn around the full perimeter of the nucleus starting at the opposite site of nucleus from where Golgi localized, and histogram measurements were obtained of fluorescence intensity along the length of the line. Radar plots were drawn based on a representative healthy and a representative patient Golgi.

To measure the volume and the mean intensities of the TGN, Z-stacks at 0.13 µm per slice were acquired and analyzed in Fiji using the “3DGolgiCharacterization” script (https://doi.org/10.5281/zenodo.10566786).

For co-localization analysis of M6PR and TGN46, and M6PR and Vps35 a rectangular roi at the perinuclear area was drawn and Mander’s coefficient was obtained using the JACoP plugin in FIJI.

To measure EEA1 size Z-stacks at 0.13 µm per slice were acquired and analyzed in Fiji using the “particleCount3D” script (https://doi.org/10.5281/zenodo.10566810).

To count LRBA^+^ endosomes in parental and Arf1+3 dKO cells, images were thresholded and a rectangle roi was drawn outside of the Golgi area and the number of endosomes were counted with analyze particles plugin using FIJI.

### Live-cell imaging of BFA and GCA inhibition

For live-cell imaging, cells were seeded on an ibidi µ-slide VI 0.4 channel slide (#80606, Ibidi) and tetracycline induced (1 µg/ml) for LRBA expression 24 hours prior to data acquisition. Live- cell imaging was performed in complete growth medium lacking phenol red at 37°C with 5% CO_2_ using an inverted Axio Observer microscope (Zeiss) with a Plan Apochromat N 63×/1.40 oil DIC M27 objective and a Photometrics Prime 95B camera. Filters with standard specifications for GFP were used to image 3xFlagEGFP-LRBA. After selecting cells to be imaged, first a snapshot was taken and then BFA (at 2 µg/ml final concentration) or GCA (at 10 µM final concentration) was applied using a BioRad Econo Pump system with 4 ml/min flow rate for immediate imaging. In case of BFA imaging, images were taken every 2 seconds for 2 minutes. For GCA, images were acquired every 2 seconds for 3 minutes then every 30 seconds for 2 minutes.

### Live-cell imaging of lysosomes using Magic Red and Lysotracker Green

For live-cell imaging, cells were seeded on an imaging chamber (ibidi µ-slide) 24 h prior to data acquisition. Magic Red (ImmunoChemistry Technologies #SKU:937) and Lysotracker Green (10 nM, Invitrogen #L7526) were diluted in imaging medium (phenol-red free CM). This staining solution was topped up with fresh imaging medium after 1 hour incubation. Live-cell imaging was performed at 37 °C with 5% CO_2_ using an inverted Axio Observer microscope (Zeiss) with a Plan Apochromat N 63×/1.40 oil DIC M27 objective and a Photometrics Prime 95B camera. Filters with standard specifications for GFP and dsRed were used to image LysoTracker Green or MagicRed signal. To measure lysosome distribution, Magic Red signal was quantified by drawing a line starting from the nucleus towards the cell periphery. Histogram measurements were obtained of fluorescence intensity along the length of the line and normalized to the minimum value of each cell which were then averaged to each cell line.

### Golgi reassembly analysis

Fibroblasts were seeded onto fibronectin coated glass coverslips and let adhere overnight. Coverslips were treated with 10 μM golgicide A or DMSO (diluted in CM) for 2 hours (2 coverslips at this point were fixed). Cells were washed 3x with PBS and incubated in CM for indicated time points. After incubation, coverslips were rinsed with PBS and fixed with 4% PFA for 10 minutes RT. Samples were washed 3x with PBS and permeabilized using 0.1% Triton in PBS for 5 minutes at RT. Coverslips were rinsed once with PBS, blocked with 5% FBS for 1 hour and incubated with anti TGN46 antibody (#AHP500GT, BioRad, 1:2000) at 4°C overnight. Coverslips were washed 3x with PBS prior to secondary antibody incubation. After 1 hour incubation with AlexaFluor568 secondary antibody, coverslips were washed 3x5min with PBS and mounted with Vectashield (Vector Laboratories, Inc. #H-1000) mounting medium containing Hoechst33342 (#639, ImmunoChemistry Technologies, 200x) and sealed with nail polish. Coverslips were imaged using the confocal Olympus Fluoview FV3000 system, an UPLSAPO 60x/1.30 objective with silicone oil.

### EGF uptake assay

Fibroblasts were seeded on fibronectin coated glass coverslips and let adhere overnight. Cells were washed 3x with PBS and incubated in serum-free DMEM for 3 hours. Cells were then washed 3x with ice-cold PBS and incubated with 2 μg/ml EGF-TexasRed (diluted in serum-free DMEM) at 4°C for 30 minutes. Then cells were washed 3 times with ice-cold PBS and endocytosis was stimulated with unlabeled 2 μg/ml EGF (diluted in serum free DMEM) at 37°C for indicated time points (0-5-15-30-45-60 minutes). Cells were then rinsed fast with ice cold PBS and fixed with 4% paraformaldehyde for 10 minutes at RT. Coverslips were washed 3 times with PBS and mounted in Vectashield containing Hoechst33342 dye to visualize nuclei and sealed with nail polish. Coverslips were imaged using the confocal Olympus Fluoview FV3000 system, an UPLSAPO 60x/1.30 objective with silicone oil. EGF uptake was quantified by measuring the fluorescence intensity of each cell.

### EGFR signaling measurements

Fibroblasts were seeded on 6 well plates and let adhere overnight. Cells were washed 3x with PBS the next day and incubated in serum-free DMEM overnight. Cells were stimulated with unlabeled 2 μg/ml EGF (diluted in serum free DMEM) at 37 °C for indicated time points (0-5-15-30-45-60 minutes). Cells were then rinsed fast with ice cold PBS and lysed using M-PER lysis buffer (supplemented with protease inhibitor (Roche) and HALT phosphatase inhibitor (Thermo Scientific^TM^ #87786) and incubated in a multi shaker for 10 minutes at 4°C. Lysates were centrifuged at 4°C and 13,000 rpm for 10 minutes. Protein concentration was determined in the supernatant using BCA assay (#23228, Thermo Fischer). Lysates were adjusted to the same concentration with lysis buffer and diluted with Laemmli. Samples were denaturated at 65°C for 10 minutes. Samples were resolved by 10% SDS-PAGE and transferred onto nitrocellulose membrane (Amersham) using wet transfer for 3 hours. Membranes were blocked with TBST (20 mM Tris, 150 mM NaCl, pH 7.6, 0.1% Tween20) with 5% non-fat dry milk for 30 min and incubated with primary antibodies, rabbit monoclonal anti EGF receptor (D38B1) XP® antibody (#4267, Cell Signaling, 1:1000), rabbit monoclonal phospho-EGF receptor (Tyr1068) (D7A5) XP® antibody (#3777, Cell Signaling, 1:1000), mouse monoclonal anti p44/42 MAPK (Erk1/2) (L334F10) antibody (#4696, Cell Signaling, 1:2000), rabbit polyclonal anti phospho-p44/42 MAPK (Erk1/2) (Thr202/Tyr204) antibody (#9101, Cell Signaling, 1:1000) overnight at 4°C. Mouse anti calnexin antibody clone 37 (#610523, BD Biosciences, 1:2000) was used as a loading control. After 2 hours incubation with HRP-conjugated secondary antibody (1:10,000; anti-mouse or anti-rabbit, Invitrogen #31430 and #31460) in TBST, chemiluminescence signals were detected using Immobilon Western HRP Substrate (Advansta) and imaged using a FusionFX (Vilber Lourmat). pEGFR and EGFR levels were measured with FIJI and normalized to calnexin loading controls (pEGFRnorm, EGFRnorm). Then pEGFRnorm values were normalized to EGFRnorm values and plotted over time. Similarly pERK and total ERK levels were measured and normalized to calnexin loading controls (pERKnorm, ERKnorm). Then pERKnorm values were normalized to ERKnorm values and plotted over time.

### Western blot analysis

Cells were lysed 24 hours after seeding in lysis buffer (1% Triton X-100, 150 mM NaCl, 20 mM Tris pH 7.5, 1mM EDTA, 1mM EGTA, protease inhibitor) and denatured in Laemmli buffer at 65 °C for 10 min. Samples were resolved by 10% SDS-PAGE and transferred onto nitrocellulose membrane (Amersham). Membranes were blocked with TBST (20 mM Tris, 150 mM NaCl, pH 7.6, 0.1% Tween20) with 5% non-fat dry milk for 30 min and incubated with primary antibody, overnight at 4°C, followed by 2 h incubation with HRP-conjugated secondary antibody (1:10000; anti-mouse or anti-rabbit, Invitrogen #31430 and #31460) in TBST. Chemiluminescence signals were detected using Immobilon Western HRP Substrate (Advansta) and imaged using a FusionFX (Vilber Lourmat).

### Total RNA isolation and qRT-PCR

Total RNA was extracted and purified from 2x10^6^ fibroblasts cells using RNeasy kit following manufacturer’s instructions. RNA was subjected to reverse-transcription using GoScript reverse transcriptase primed with a mix of Oligo(dT)s (Promega) and random hexamers (Promega). qRT- PCR was performed using GoTaq qPCR master mix (Promega) and primers specific for LRBA (forward: 5’-CCAACTTCAGAGATTTGTCCAAGC-3’; reverse: 5’- ATGCTGCTCTTTTTGGGTTCAG3-’) (Wang et al., 2004), MAB21L2 (forward: 5’- CCAGGTGGAAAACGAGAGTG-3’; reverse: 5’-GGTAGAGCACCACCTCAAATTC-3’) (Deml et al., 2015), GAPDH (forward: 5’-TCAAGGCTGAGAACGGGAAG-3’, reverse; 5’- CGCCCCACTTGATTTTGGAG3-’) (Dahn et al., 2021).

### Alphafold modeling

To evaluate the structure of LRBA we used alphafold monomer v2.2 developed by Deepmind (Evans et al., 2021; Jumper et al., 2021). Due to the size of LRBA we had to create three separate models, AA1-1566, AA1000-2000 and AA1567-2863. These models were stitched together using ChimeraX according to the highest reliability values (pLDDT) of each part. To evaluate the interaction between Arf1 or Arf3 and LRBA we used alphafold multimer v2.2 (AF-M) developed by Deepmind. We had to run Arf1 or 3 with either AA1-1566 or AA1567-2863 of LRBA due to size limitations. We ran all 5 of AF-M models for 3 recycles with 3 seeding points resulting using in 15 models per run. The models were evaluated using ChimeraX.

### Stimulation of lymphocytes for functional analyses

For functional analyses of CD8^+^ lymphocytes, 2x10^5^ cells were resuspended in RPMI plus 10% FCS and cultured on a 96-well plate. Anti-CD107a-PB (#B13978, Beckman Coulter) antibody and monensin (Golgi Stop^TM^, prediluted 1:10 with RPMI + 10% FCS) were added to all conditions. Stimulation included phorbol-12-myristat-13-acetat (PMA, Sigma-Aldrich; #P8139- 5M6, concentration of 2ng/µl) and ionomycin (Sigma-Aldrich; #I0634-1M6 0,1µg/µl) for 4 hours at 37°C and 5% CO_2_. NK cell degranulation was analyzed according to an adapted protocol described by Bryceson et al. (2012) (Bryceson et al., 2012). 2x10^6^ PBMCs were either stimulated with IL-2 (Novartis, #1447583) at a concentration of 600U/ml on a 24-well plate or kept in medium. After 16 hours a manual cell analysis was performed using a Neubauer counting chamber (Optik Labor, Friedrichsdorf, Germany) and 200.000 cells were transferred on a 96-well plate. An anti-CD107a-PB antibody and monensin were added to all samples prior to stimulation. IL-2 stimulated cells were co-cultivated with K562 (#89121497) cells at a ratio 1:2 for three hours at 37°C and 5% CO_2_.

### Surface phenotyping and flow cytometry

Cells were washed in DPBS plus 0.5% human serum albumin and incubated with anti-CD45-KrO (#B36294, Beckman Coulter), anti-CD3-APC (#IM2467, Beckman Coulter), anti-CD8-APC- AF700 (#B49181, Beckman Coulter), anti-CD69-PE (#IM1943U, Beckman Coulter) and CD56- PC7 (#A21692, Beckman Coulter) for 15 minutes at RT. Cells were washed once, diluted in 300µl DPBS/HSA and measured using a ten-color flow cytometer (Navios, Beckman Coulter, Krefeld, Germany).

FACS-data were analyzed using Kaluza® version 2.1 (Beckman Coulter, Krefeld, Germany). The gating strategy is described in Figure S3 and S4. For NK cells, CD107a^+^ gates were set after a healthy and unstimulated control for each experiment, given the population as percentage of total NK cells. For CD 8^+^ T cells, the mean fluorescence intensity (MFI) was determined as an indicator of CD107a expression.

### Statistics

Statistical analysis was performed using GraphPad Prism 10.1.1 (GraphPad Software, Inc., San Diego, CA, USA). For statistical evaluation, the normality of the data was routinely tested using the Shapiro-Wilk or Kolmogorov-Smirnov normality test. For the comparison of two groups, a t test was employed, whereas to compare three or more groups, a one-way ANOVA was performed followed by a Kruskal-Wallis test.

## Acknowledgements

We are grateful to the patients, their families, and healthy donors. We thank S. Erben (Staff Labor for SCT, Frankfurt) for technical assistance, Dr. E. Jacobsen (University Hospital Ulm) for suggestions regarding CTL degranulation analysis, and the pediatric surgery for skin biopsy. The HeLa Trex FlpIn-3xFLAG-EGFP-LRBA cell line was kindly provided by Dr Serhiy Pankiv and Prof. Anne G. Simonsen (University of Oslo, Norway). We acknowledge the support of the FACS facility of the Biozentrum, Cinzia Tiberi Schmidt from the BioEM facility and IMCF facility of the Biozentrum, especially Laurent Guérard and Alexia Loynton-Ferrand. We thank Dominik Buser for helpful discussions and for antibodies. Calculations were performed at sciCORE (http://scicore.unibas.ch/) scientific computing center at University of Basel. We would like to thank Ludovic Enkler for critical reading of the manuscript.

## Contribution information

VS, LML, SB, AS designed the experiments, VS and LML performed experiments. VS, LML, DO, SB, AS, analyzed data, AS, SB, MH supervised the study, NZ provided patient material and VS, AS and SB wrote the manuscript with input from all authors.

## Supplementary figure legends

**Supplementary figure 1.**
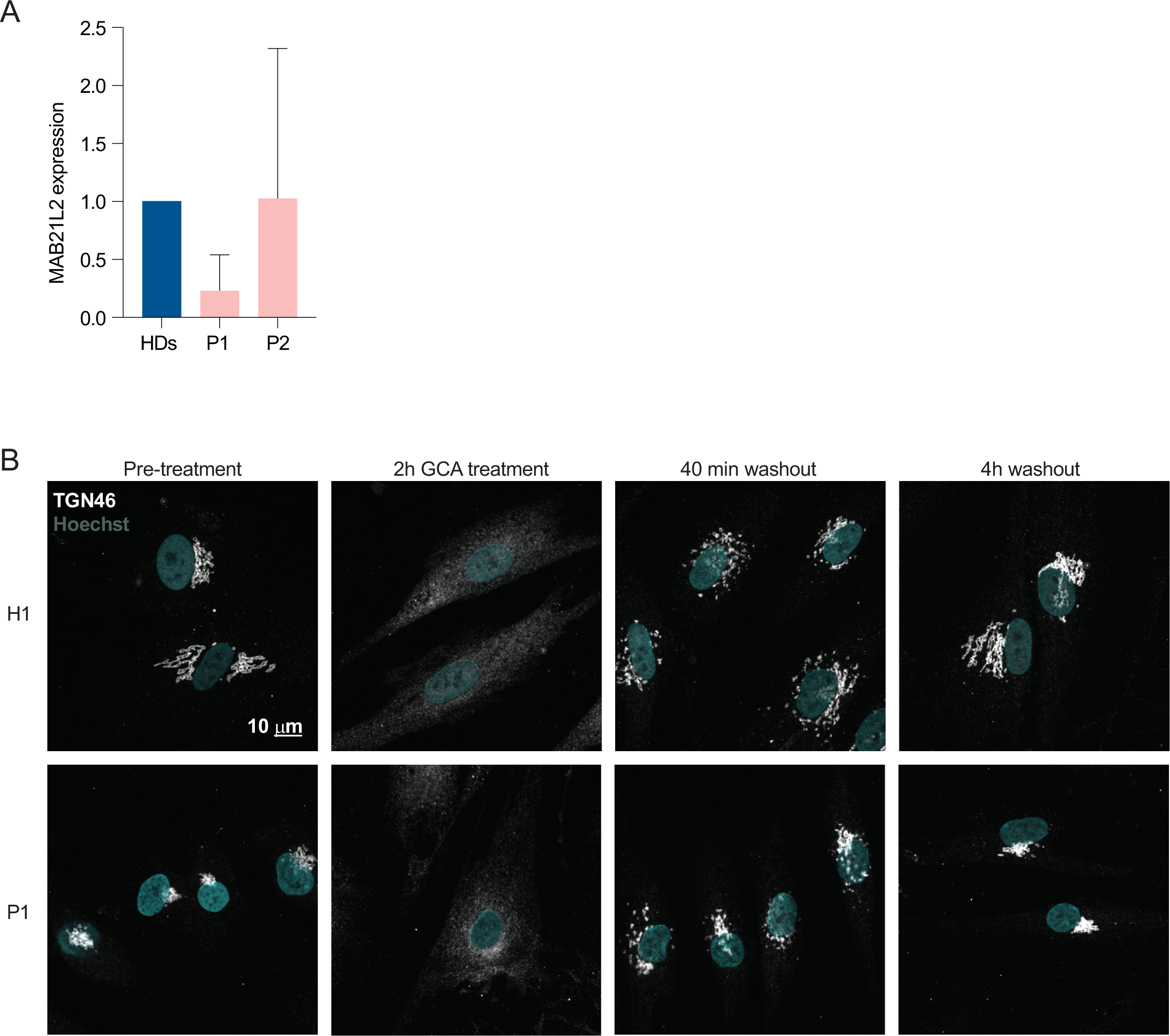
LRBA does not regulate Golgi assembly. (A) The nested MAB21L2 gene expression levels in LRBA deficient fibroblasts. MAB21L2 mRNA levels in two patients and three HD fibroblasts were determined by qRT-PCR. Mean and standard deviation are shown from n=3 biological replicates. **(B)** Golgi reassembly is not regulated by LRBA. H1 and P1 cells were grown on coverslips and treated with GCA for 2 hours to vesiculate the Golgi. After 2 hours GCA was washed out and the cells were incubated with complete growth media for indicated time points for Golgi reassembly. Cells were then fixed and their Golgi was visualized by endogenous TGN46 staining and their nuclei by Hoechst staining. Representative maximum Z-projected images are shown.

**Supplementary figure 2.**
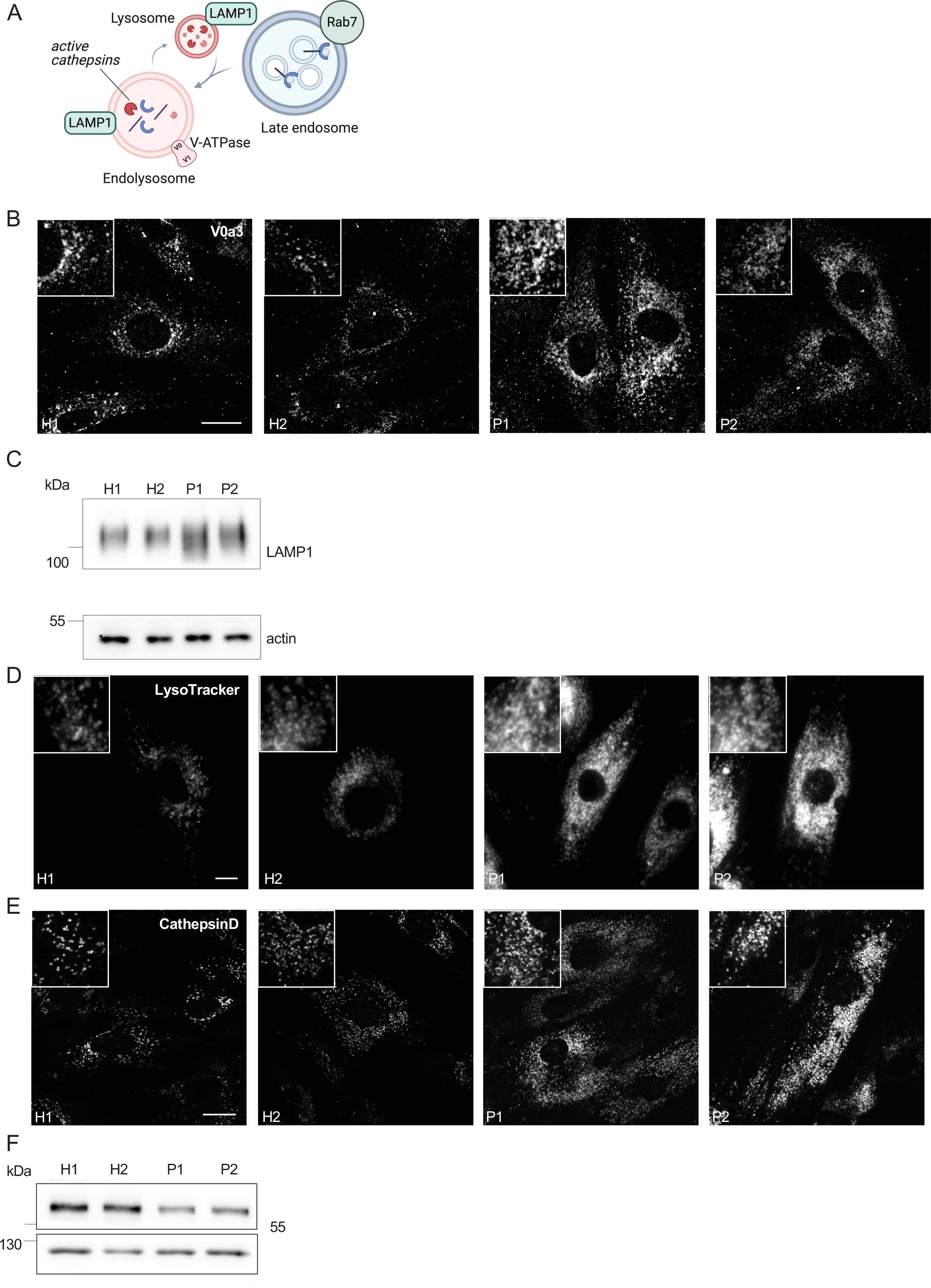
Enlarged (endo)lysosomal structures are acidified and contain active cathepsin D. (A) Scheme showing Rab7^+^ late endosome fusing with LAMP1+ lysosomes and becoming endolysosomes. Endolysosomes are acidified, contain active cathepsins in their lumen and vacuolar-ATPase and LAMP1 in their membrane. **(B)** Visualization of endolysosomes by immunostaining the V0 subunit of the V-ATPase. Healthy and LRBA deficient fibroblasts were fixed and stained with V0a3 antibody. Representative confocal images. Inlays are shown in the top left corner of the images. Scale bar, 10 μm. **(C)** Immunoblot analysis of LAMP1 protein levels in 2 healthy and 2 LRBA patients’ fibroblasts. Actin was used as a loading control. **(D)** Accumulation of acidified endolysosomes in LRBA deficient fibroblasts. Fibroblasts were seeded onto imaging chambers, stained with LysoTracker Green and imaged live at 37°C and 5% CO_2_ atmosphere. Maximum Z-projection of wide-field images are shown. Inlays are shown in the top left corner of the images. Scale bar, 10 μm. **(E)** Cathepsin D is present in accumulating, enlarged (endo)lysosomes. Healthy and LRBA deficient fibroblasts were fixed and stained with endogenous cathepsin D antibody. Representative confocal images. Inlays are shown in the top left corner of the images. Scale bar, 10 μm. **(F)** EGFR protein levels are reduced in LRBA deficient fibroblasts. Immunoblot analysis of EGFR in 2 healthy and 2 LRBA patient fibroblast cell lines. Actin was used as a loading control.

**Supplementary figure 3.**
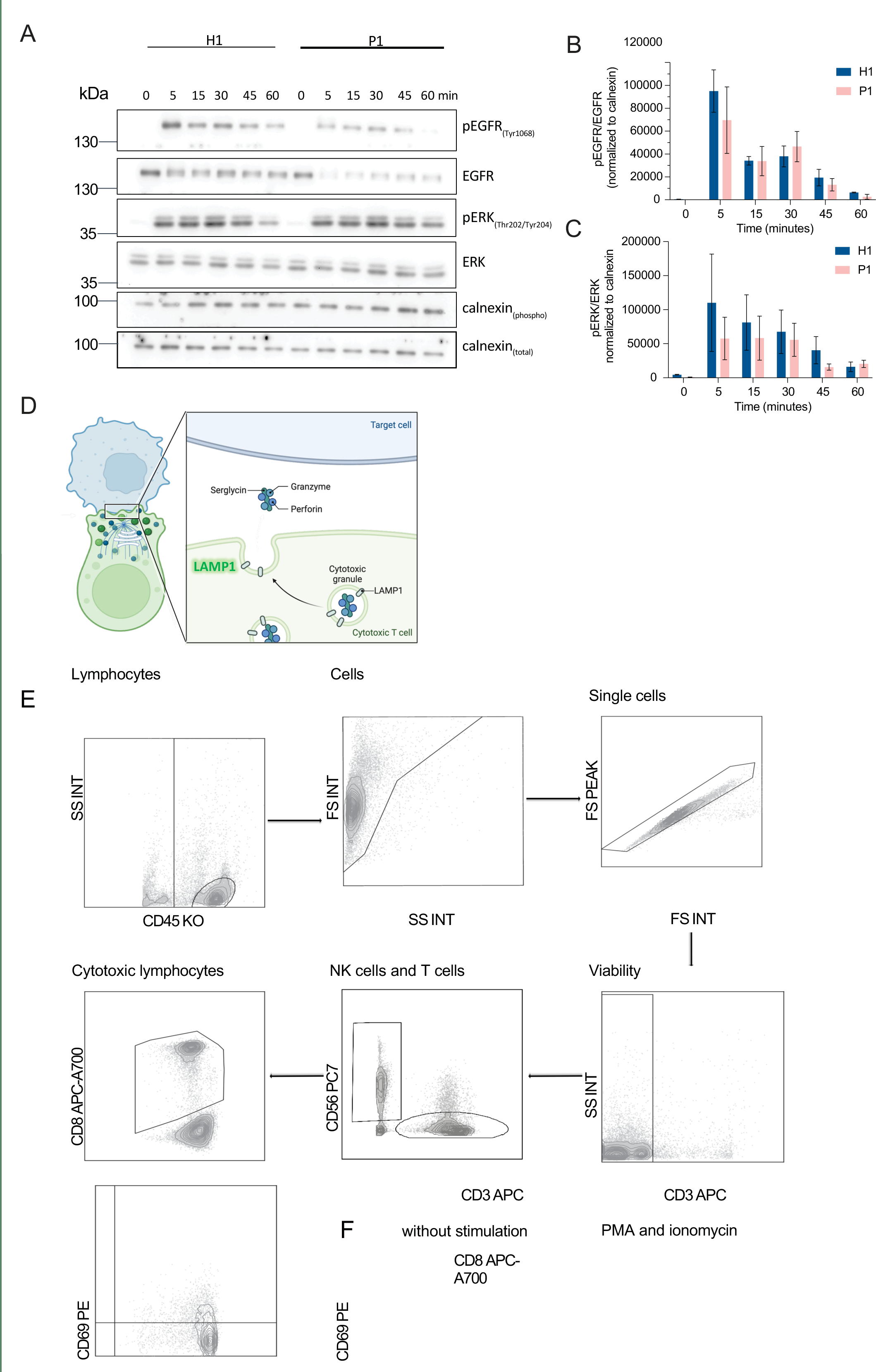
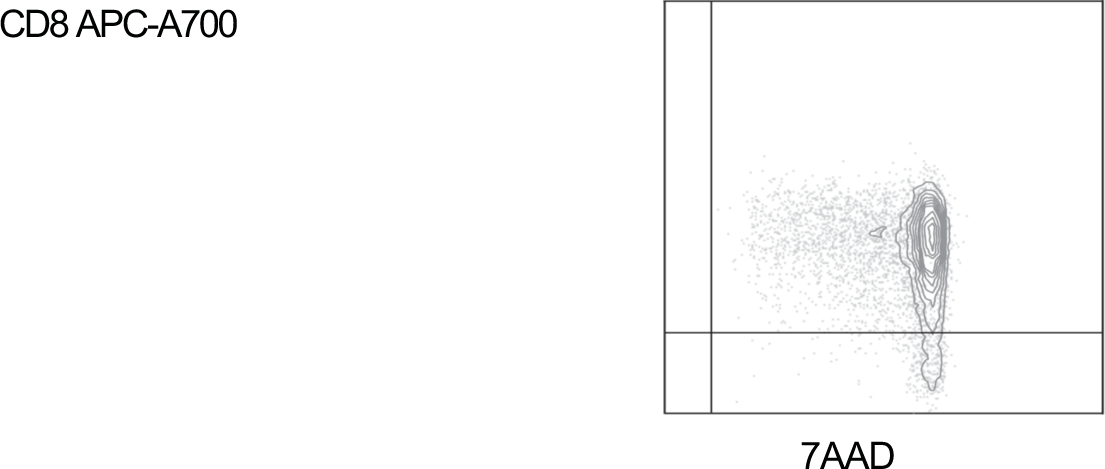
LRBA does not regulate EGFR signaling attenuation. (A) Immunoblot analysis of EGFR signaling kinetics in H1 and P1 fibroblast cell lines. Cells were serum starved overnight in DMEM and were then incubated with 2 μg/ml EGF in serum-free DMEM for indicated time points. Cells were then rinsed with ice-cold PBS and lysed with M-PER lysis buffer supplemented with protease and phosphatase inhibitor. For addressing EGFR signaling, antibody against EGFR and its phosphorylation site Tyr1068 was used. We also detected the downstream ERK phosphorylation with the Thyr202/204 phosphorylation site specific antibody. Calnexin was used as loading control. **(B)** Quantification of pEGFR levels and kinetics upon EGF stimulation based on immunoblots shown in (A). pEGFR and EGFR levels were measured and normalized to calnexin loading controls (pEGFR_norm_, EGFR_norm_). Then pEGFR_norm_ values were normalized to EGFR_norm_ values and plotted over time. **(C)** Quantification of pERK levels and kinetics upon EGF stimulation based on immunoblots as shown in (A). pERK and total ERK levels were measured and normalized to calnexin loading controls (pERK_norm_, ERK_norm_). Then pERK_norm_ values were normalized to ERK_norm_ values and plotted over time. **(D)** Scheme of immunological synapse formation and lytic granule exocytosis in cytotoxic T cells. Upon target cell recognition immune synapse is formed which induces a strong polarization in T cells. The MTOC is trafficked to the immunological synapse bringing other organelles like the Golgi network, endosomes and lytic granules to the synapse. Lytic granules are lysosome-related organelles containing canonical lysosomal proteins (LAMP1), granzyme and perforin. During degranulation, the lytic granules are fused with the plasma membrane. Upon release, perforin mediates generation of pores in the plasma membrane of the target cell allowing granzymes to access the cytoplasm and induce apoptosis. The degranulation process exposes LAMP1 on the cell surface and can be used as a marker for degranulating cells. **(E)** Gating strategy of CD8+ T cells and NK cells in degranulation assay. Gating strategy for cytotoxic lymphocytes and NK cells is shown on the sample of a HD. Leukocytes and lymphocytes are definied in SSC/CD45 gate. Lymphocyte populations are visualized again in a SSC/FSC gate. Single cells are discriminated from doublets in a FS peak/FSC gate. Viable cells are gated as 7AAD negative cells in a viability gate. NK cells are defined as CD3^-^ and CD56^+^ cells. Cytotoxic lymphocytes are defined as CD3^+^ and CD8^+^. **(F)** CD69 was used as a common marker for lymphocyte activation upon stimulation. An example of CD69 gating in CD8^+^ cells from a HD is shown. SSC, side scatter, FSC, forward scatter.

**Supplementary figure 4.**
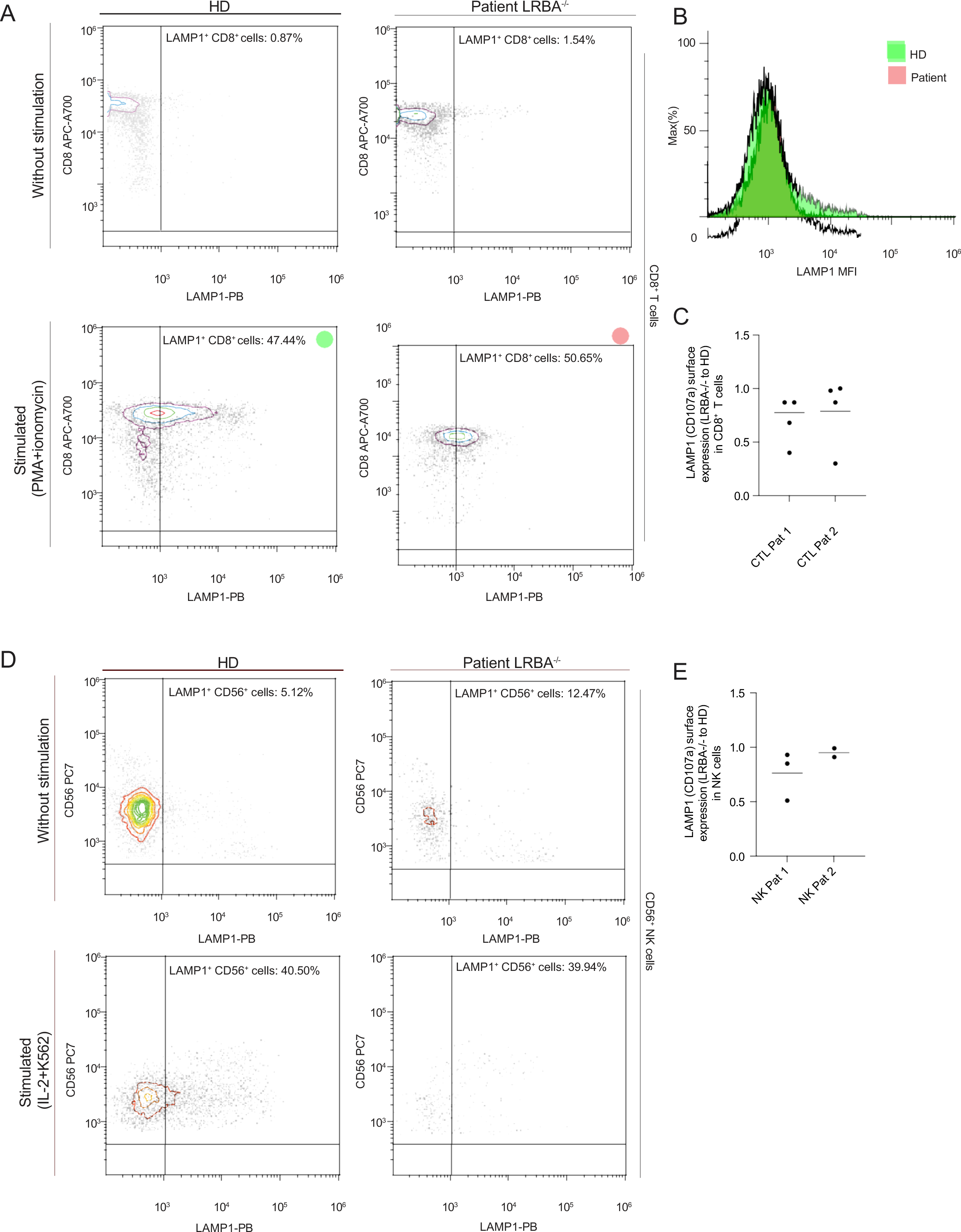
**Degranulation of CD8+ T cells and NK cells is unimpaired in LRBA deficiency**. **(A)** Flowcytometric analysis of LAMP1(CD107a) surface expression in CD8^+^ T cells of healthy individual (HD) and LRBA deficient patient (LRBA^-/-^) in mononuclear cell (PMCS) isolated from peripheral blood samples. Upon stimulation with PMA+ionomycin there is a significant increase in LAMP1 expression. Dot blots are showing the CD8^+^LAMP1^+^ population marked with a green circle for the healthy and red circle for the affected individual. **(B)** The mean fluorescence intensity (MFI) was calculated for each sample, given as green (HD) and red (patient) histograms. **(C)** In repetitive analyses, patient cells degranulate almost as efficient as healthy cells. Degranulation is given by the ratio of LAMP1 surface expression for each LRBA deficient patient and the healthy control in each degranulation assay as these where available at different time points. **(D)** For the NK cells the stimulation included IL-2 and co-culture with K562 cells, resulting in a subpopulation of NK cells (%) expressing LAMP1 in healthy donor and patient. Dot bolts show the percentage of the CD56^+^LAMP1^+^ population (top right quadrants). **(E)** Repetitive analyses for each patient show that patients 1 has a higher variability than patient 2. Both are capable of degranulating LAMP1^+^ lytic granules.

**Supplementary figure 5.**
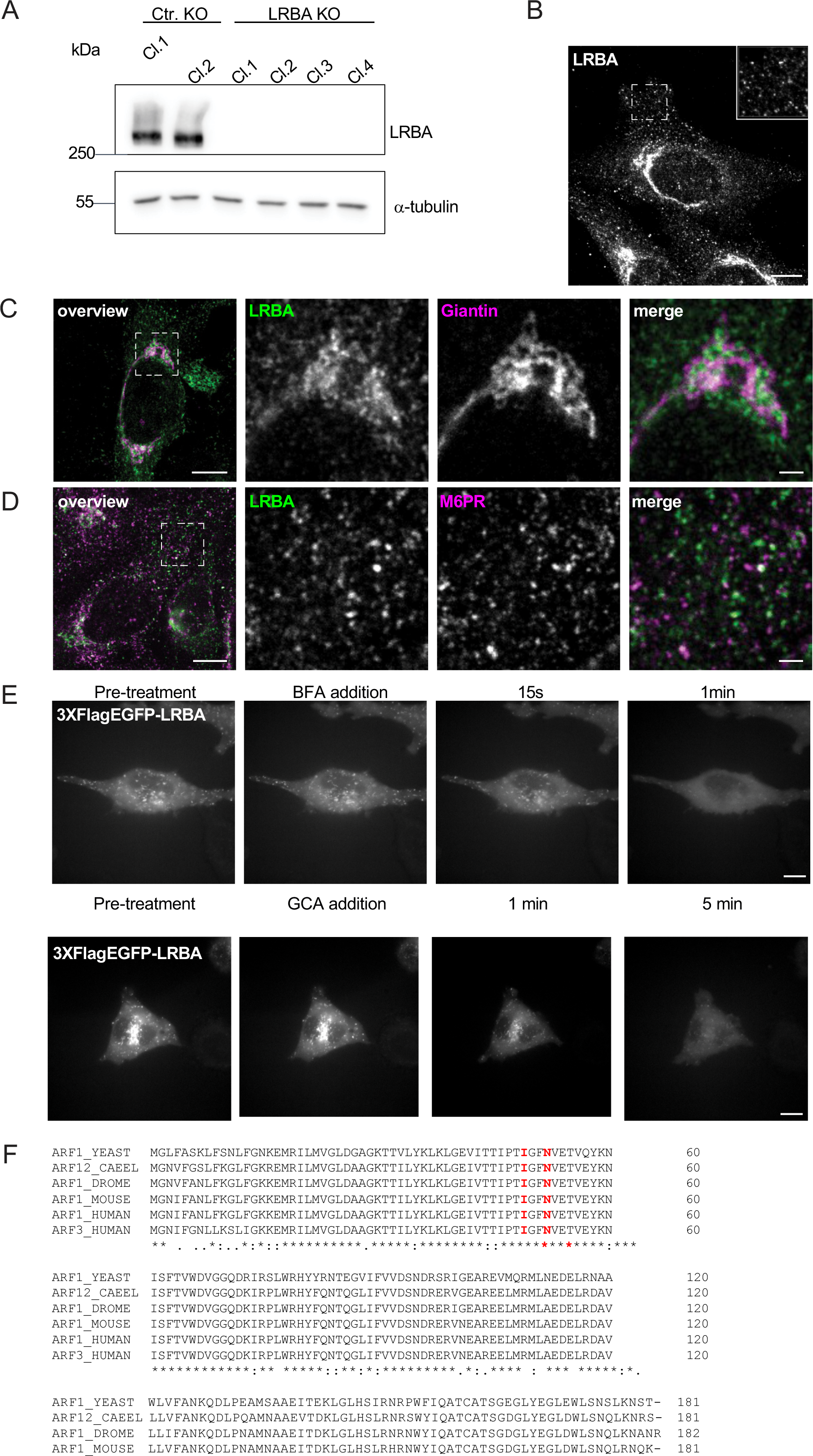
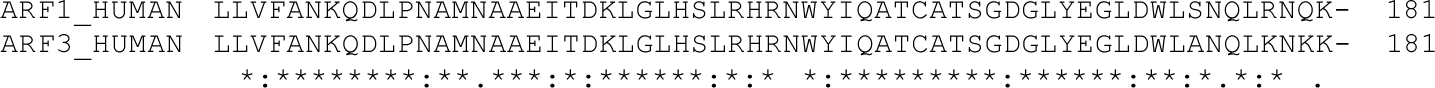
LRBA is endogenously expressed in HeLa cells and recruited to endosomes by Arfs. (A) Immunoblot analysis of LRBA presence in HeLa cells of 2 control and 4 LRBA KO clones using polyclonal LRBA antibody and *α*-tubulin as a loading control. **(B)** LRBA is localized at the perinuclear region and on vesicular structures in HeLa cells. Immunofluorescence analysis of endogenous LRBA in fixed HeLa cells. Scale bar, 10 μm. **(C)** LRBA appears juxtaposed to the cis-Golgi in HeLa cells. Immunoblot analysis of endogenous LRBA and the cis-Golgi marker giantin colocalization in fixed HeLa cells. Squares show magnification of the perinuclear area. The labeling of the single channels represents the color of the channel on the merged image. Scale bar, 10 μm, inlays 2 μm. **(D)** LRBA does not colocalize with M6PR in HeLa cells. Immunoblot analysis of endogenous LRBA and endogenous M6PR colocalization in fixed HeLa cells. Squares show magnification of the perinuclear area. The labeling of the single channels represents the color of the channel on the merged image. Scale bar, 10 μm, inlays 2 μm. **(E)** LRBA dissociates from membranes upon ArfGEF inhibitors. Live-cell imaging of 3xFlagEGFP-LRBA upon BFA (top panels) and GCA (lower panels) inhibition. Scale bar, 10 μm. **(F)** The amino acids isoleucine 49 and the aspargine 52 of Arf1 and Arf3 were predicted to interact with LRBA. Both amino acids are conserved across species. The amino acid sequences of the yeast, *C.elegans* (CAEEL), *Drosophila melanogaster* (DROME), mouse and human Arf1 and human Arf3 were aligned. Labels: (*) conserved sequence; (:) conservative mutation; (.) semi-conservative mutation; (-) gap.

